# Integrated single-cell multiomic analysis of HIV latency reversal reveals novel regulators of viral reactivation

**DOI:** 10.1101/2022.07.26.501558

**Authors:** Ashokkumar Manickam, Jackson J Peterson, Yuriko Harigaya, David M Murdoch, David M Margolis, Alex Oesterling, Zhicheng Guo, Cynthia D Rudin, Yuchao Jiang, Edward P Browne

## Abstract

Despite the success of antiretroviral therapy, HIV cannot be cured because of a reservoir of latently infected cells that evades therapy. To understand the mechanisms of HIV latency, we employed an integrated single-cell RNA-seq/ATAC-seq approach to simultaneously profile the transcriptomic and epigenomic characteristics of ~4000 latently infected cells after reactivation using three different latency-reversing agents (LRAs). Differentially expressed genes and differentially accessible motifs were used to examine transcriptional pathways and transcription factor (TF) activities across the cell population. We identify cellular transcripts and TFs whose expression/activity was correlated with viral reactivation and demonstrate that a machine learning model trained on these data was 68% accurate at predicting viral reactivation. Finally, we validate the role of a new candidate HIV-regulating factor, GATA3, in the viral response to prostratin stimulation. These data demonstrate the power of integrated multimodal single-cell analysis to uncover novel relationships between host cell factors and HIV latency.

## Introduction

The formation of a latently infected reservoir in CD4 T cells is a key barrier to an HIV cure (Chun et al., 1997; Finzi et al., 1997). The reservoir is highly stable, long lived, and resistant to current antiretroviral therapy (Brodin et al., 2016; Elsheikh et al., 2019; Ismail et al., 2021). Additionally, clonal expansion in vivo counteracts gradual erosion of infected cells (De Scheerder et al., 2019; Lau et al., 2021). A key mechanism in the formation of this reservoir is the phenomenon of viral latency, in which HIV transcription is reversibly silenced post integration, allowing it to evade the host immune defenses. Sporadic reactivation of these cells generates ‘blips’ of viremia during therapy (Crowell et al., 2020; Rong and Perelson, 2009) and seeds rapid rebound upon interruption of therapy (De Scheerder et al., 2019). Current cure strategies for HIV involve transiently inducing viral gene expression in latent proviruses using latency-reversing agents (LRAs), followed by immune clearance of the reactivated cells. However, this approach has thus far only achieved limited success. Reactivation of latently infected cells with existing LRAs is inefficient, with typically ~10% of replication-competent proviruses being reactivated in patient-derived cells, even with ‘potent’ LRAs (Ho et al., 2013). For the LRA/clearance approach to be successful, broad reactivation of the reservoir will be required. This inefficiency of latency reversal with LRAs is likely due to a combination of stochastic processes that regulate viral gene expression, the existence of multiple layers of repression to HIV gene expression, and heterogeneity in both the cellular environment and proviral integration sites in infected cells. Fully defining the nature of these repressive mechanisms is urgently needed to allow the development of broader acting LRAs or approaches with a combination of different LRAs. Prior work indicates that silencing of HIV in CD4 T cells involves the combined effects of low levels of positive TFs, such as NF-kB and AP1 (Hokello et al., 2021; Wong and Jiang, 2021), active repression by cellular factors (e.g., NELF, DSIF) (Ait-Ammar et al., 2019), sequestration of the elongation complex pTEFB (CyclinT1/CDK9) (Mbonye et al., 2021), and histone modifications that create a repressive heterochromatin environment around the virus promoter (Turner and Margolis, 2017; Zhang et al., 2017). Nevertheless, the regulation of HIV gene expression remains incompletely understood, and additional mechanisms likely exist that will need to be fully characterized.

Gene expression is regulated by transcription factors (TFs) that bind to promoter or enhancer regions near the gene, and mediate chromatin remodeling of these regions (Li et al., 2015), during which nucleosomes are removed, repositioned, or modified by covalent molecules such as acetylation or methylation (Lorch et al., 2006). This remodeling then allows recruitment of additional TFs, as well as RNA polymerase II and transcriptional elongation factors to promoters and enhancers (Quevedo et al., 2019; Stees et al., 2012). Thus, TF binding, chromatin remodeling, and RNA transcription are key regulatory factors in cellular gene expression that act in a coordinated fashion to determine the expression level for each gene. However, fully understanding how these processes interact to regulate gene activity in individual cells has been difficult due to the technical limitations of profiling individual cells. Recent advances in multimodal single cell analyses now allow concurrent observation of both the RNA levels for a given gene and the accessibility of the chromatin associated with this gene in single cells. These approaches have led to fundamental new insights into the regulation of gene expression.

Thus, in this study, we used an integrated single-cell RNA sequencing (scRNA-seq) and single-cell sequencing assay for transposase-accessible chromatin (scATAC-seq) approach to characterize transcriptomic and epigenomic profiles from the same cells in a cell line model of HIV latency to uncover the networks of transcriptional regulators that control HIV gene expression in response to hallmark LRAs. By correlating HIV RNA levels with individual cellular transcripts, as well as with chromatin peak dynamics and TF activity scores inferred from ATAC-seq peak accessibility, we revealed several novel candidate regulators of HIV gene expression. Finally, by using shRNA knockdown, we identify GATA3 as a novel regulator of HIV latency and reactivation.

## Results

### Latency reversal in a cell line model of HIV latency

To understand the relationship between cellular transcriptome, chromatin state, and HIV transcription, we stimulated 2D10 cells (Pearson et al., 2008), a cell line model of HIV latency, with one of three different latency-reversing agents (LRAs), or with control vehicle (DMSO) (**Figure 1A**). The LRAs we used represented different mechanisms of action - vorinostat (HDACi), iBET151 (BRDi), and prostratin (PKCa). We added limiting amounts of each LRA to induce HIV reactivation in only a subset of cells (20-30%) at 24h post stimulation (**Figure 1B**). In the presence of control vehicle only, approximately 7% of the cells displayed some level of background HIV expression as measured by detection of Green Fluorescent Protein (GFP). Each population was then analyzed by integrated scRNA-seq/scATAC-seq, yielding data from ~1000 cells for each condition, for a total of ~4000 cells. Quality control metrics indicated high-quality library preparation and sequencing quality with well-balanced depth of coverage between the ATAC and RNA domains (**Figure S1**). Dimension reduction using Uniform Manifold Approximation and Projection (UMAP) (Becht et al., 2018) of the scRNA-seq data showed a clear separation of LRA treated cells compared to the control (DMSO) treated cells along UMAP1, indicating LRA-specific modulation of cellular genes (**Figure 1C, left panel**). Additionally, both DMSO and LRA-treated cells could roughly be separated into two large groupings separated along UMAP2. By contrast, UMAP display of the matching scATAC-seq data indicated that separation of the four conditions at the epigenomic level was less clear. DMSO and iBET151 occupied overlapping regions, indicating that iBET151 causes limited rearrangement of overall chromatin accessibility. Vorinostat (VOR) and prostratin-treated cells were somewhat separated from other cell conditions within the scATAC-seq UMAP projection but were still part of an overall closely connected single cluster of cells (**Figure 1C, right panel**). This lower degree of separation of the cells from different conditions using the ATAC-seq data likely reflects the sparse nature of scATAC-seq data as well as possibly a lower level of epigenomic diversity compared to transcriptomic diversity in the population of cells. We found that this observation was true when the cells were clustered using four different metrics derived from the ATAC-seq data (peak count, peak count with additional genomic annotations and external regulomic data, gene accessibility, and TF motif deviation) (**Figure S2**). We also labeled ten individual clusters of cells (clusters 0-9) based on their transcriptome, irrespective of condition, using graph-based clustering (**Figure 1D**). This clustering pattern indicated that diverse states of the cells exist within each condition. Similar to our observation when the cells were labeled by condition, we observed that these transcriptomic clusters were harder to discern from the scATAC-seq data.

**Figure 1.**
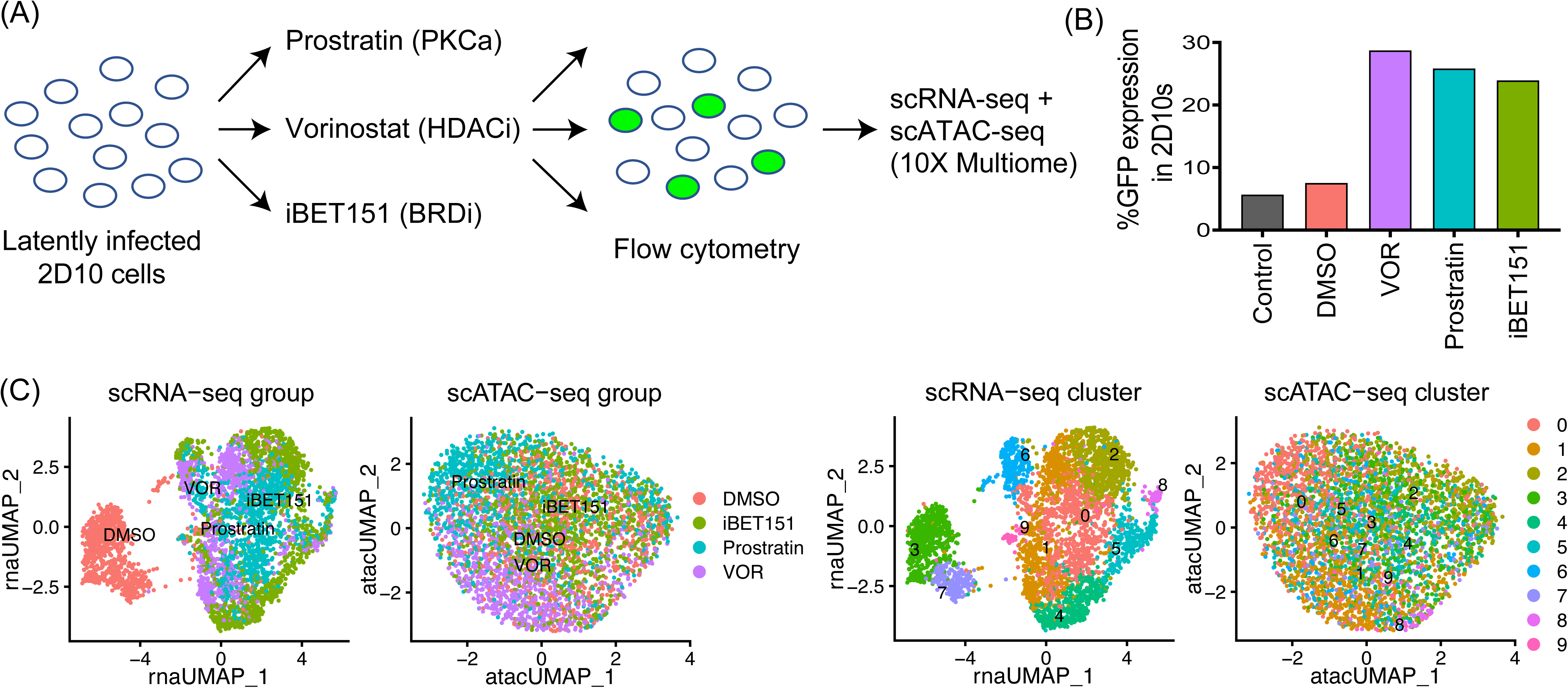
Single cell multiomic analysis of HIV latency reversal. **(A)** A schematic of the experimental design of HIV latency reversal followed by scRNA-seq and scATAC-seq multiomic sequencing. **(B)** Percentage of cells expressing the GFP reporter in the control (DMSO), as well as in those treated with LRAs (Vorinostat (VOR), prostratin, and iBET151). **(C)** UMAP dimension reduction of scRNA-seq and scATAC-seq, with cells labeled by conditions. **(D)** UMAP dimension reduction of scRNA-seq and scATAC-seq, with cells labeled by cluster.

### Heterogeneous viral gene expression in individual cells

We next examined the expression of HIV transcripts across the cell population. Reads were aligned to an HIV reference genome derived from the viral clone used to generate the 2D10 cells (Kim et al., 2011). As expected, each LRA stimulation caused a significant increase in the abundance of HIV-mapping reads in the cells, with significant *p*-values from the Turkey test and one-way analysis of variance (ANOVA) test (**Figure 2A**). Specifically, 20.0 percent of cells had detectable viral RNAs (vRNAs) with at least one unique molecular identifier (UMI) read for DMSO treated cells, whereas the percentages were 41.0, 52.0, and 43.0 upon treatment with iBET151, prostratin, and VOR, respectively. The higher fraction of cells with detectable vRNA compared to the GFP+ fraction observed by flow cytometry indicates that flow cytometry for GFP underestimates the fraction of cells with active viral gene expression. Abundance of vRNA reads across the vRNA+ cells within each condition was highly variable and ranged from 0.6 to 17.3 percent of the reads for each cell. Interestingly, we observed that the average expression of vRNA across the population was significantly higher for VOR-treated cells than for the other LRAs, despite a similar percentage of cells upregulating vRNA and becoming GFP+ (**Figure 2A, 2B**, **Figure S3A**). This could reflect additional post-transcriptional blocks to viral expression in VOR treated cells that are not present in the other conditions. That is, at these concentrations, VOR more efficiently reactivates transcription in responding cells, but protein level expression is inhibited by additional blocks that are overcome more effectively by prostratin or iBET151. Alternatively, VOR may itself trigger a post-transcriptional block to expression not present with other LRAs.

**Figure 2.**
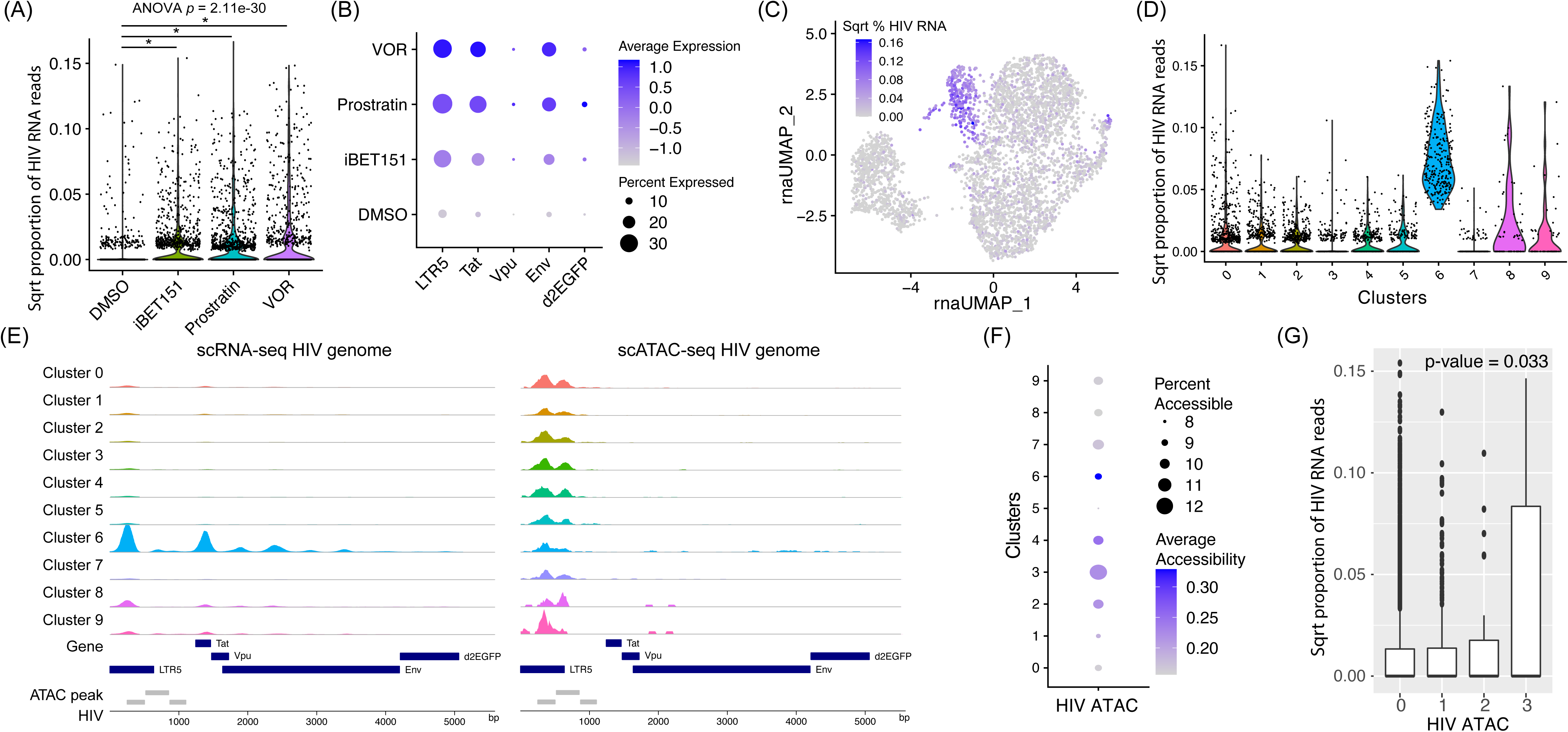
HIV viral RNA expression and chromatin accessibility. **(A)** Transformed HIV RNA reads under different treatment conditions. *p*-value from one-way ANOVA test is shown. The asterisks represent significant pairwise differences based on the Turkey test (*p* = 3.25e-8 between iBET151 and DMSO, *p* = 1.93e8 between Prostratin and DMSO, and *p* = 1.93e-8 between vorinostat and DMSO). **(B)** Dot plots of RNA transcript levels for the regions of the HIV genome: The long terminal repeat (LTR), Tat, Vpu, Env, and d2EGFP. **(C)** UMAP plot of the scRNA-seq data color-coded based on the levels of HIV RNA expressions. **(D)** Transformed total HIV RNA reads across different cell clusters. **(E)** Coverage plots showing normalized read counts from scRNA-seq (left) and scATAC-seq (right) along the HIV genome for different cell clusters. The dark blue and gray boxes represent gene annotations and ATAC peaks. **(F)** Dot plot showing ATAC-seq signals summed across the entire HIV genome for different clusters. **(G)** HIV RNA expression is marginally correlated with HIV chromatin accessibilities. The bar plot shows the squared proportion of HIV RNA reads, stratified by the total number of HIV ATAC read counts, which has a maximum of three due to the sparsity of scATAC-seq. The *p*-value was obtained from a Pearson correlation testing.

Although the molecular methodology to generate the scRNA-seq libraries involved oligo-dT mediated reverse transcription, and was thus expected to generate vRNA mapping reads mainly from the long terminal repeat (LTR) of the viral genome adjacent to the HIV polyA site, we were nonetheless able to detect abundant reads from other parts of the viral genome, particularly from the Tat region (**Figure 2B, Figure S4**). This phenomenon is likely due to internal priming of cDNA synthesis in A-rich viral sequences within the viral genome. Thus, we also examined the abundance of reads that mapped to individual viral transcripts expressed by this reporter virus (Tat, Rev, Vpu, Envelope), the dreGFP reporter gene (located within the Nef transcript), as well as reads that map to the viral LTR (**Figure 2B**). Due to the identical sequences in the 5’ and 3’ LTR, all reads that mapped within the LTR region were aggregated and considered as a single region. At baseline (DMSO) only a small fraction of cells expressed RNAs that mapped to any of these regions (0.4-13.4%). After LRA stimulation, reads that mapped to the LTR, Envelope and Tat were abundantly induced for all LRA conditions, while reads mapping to Vpu, and dreGFP were less upregulated. Consistent with our previous observations, VOR induced a similar fraction of vRNA+ responding cells to the other LRAs, but a significantly higher average level of vRNA expression on a per-cell basis (**Figure 2B, Figure S4**).

We next examined whether induction of viral gene expression was linked to the transcriptomic or epigenomic characteristics of the infected cells. UMAP display of the scRNA-seq and scATAC-seq data showed that within the scRNA-seq data, vRNA expressing cells were strikingly concentrated in a defined region of the plot (**Figure 2C**). Notably, we observed that the cells that expressed high vRNA were concentrated in cluster 6 (**Figure 2D, 2E**). This clustering of vRNA+ cells was less prominent when vRNAs were excluded from the dataset when performing clustering, indicating that the viral transcripts themselves were important contributor to the formation of cluster 6 (**Figure S5**). Nevertheless, vRNA+ cells displayed a non-random distribution across the scRNA-seq UMAP plot even without inclusion of vRNA, indicating that reactivation of viral gene expression was correlated with cellular pathways and/or the induction of viral gene expression generated a distinct transcriptional response by the host-cell transcriptome.

### Viral gene expression is correlated with increased accessibility of the viral genome

To understand how the epigenomic configuration of the viral genome was related to viral genome expression, we next examined the accessibility of the HIV provirus within our dataset by examining scATAC-seq reads that mapped to the HIV genome. Overall, the HIV-mapping ATAC-seq data was extremely sparse, with 80% of cells exhibiting no detectable HIV mapping ATAC-seq reads (**Figure 2F**). Of the 20% of cells that did contain HIV mapping reads, the majority contained a single HIV read, while only a small number of cells contained two or three reads. Notably, we observed a modest but statistically significant (*p*-value = 0.033) correlation between HIV mapping ATAC-seq reads and the abundance of vRNA within each cell (**Figure 2G**), consistent with the hypothesis that viral gene expression is linked to overall proviral chromatin accessibility.

By aggregating scATAC-seq data for all cells within each condition, we were able to observe three accessibility peaks within the HIV genome (**Figure 2E**). The three peaks are all located at the 5’ end of the viral genome, overlapping with the 5’ LTR and the 5’ end of the Gag gene. The HIV genome contains several known nucleosomal positions within the 5’ end of the virus – (Nuc0 Nuc1, Nuc2, and Nuc3) (Demarchi et al., 1993; Van Lint et al., 1996; Verdin, 1991; Verdin et al., 1993). Peak 1 (HIV: 251-550) roughly corresponds to a known DNase1 hypersensitive (DHS) region between Nuc0 and Nuc1 and overlaps the viral transcription start site (TSS) at nucleotide 455. Peak 2 (HIV: 555-850) is immediately downstream of Nuc1 between Nuc1 and Nuc2, while peak 3 (HIV: 851-1100) roughly overlaps the location of Nuc3. Interestingly, of these peaks, only peak 3 displayed a consistent increase in accessibility after LRA stimulation (**Figure S6**). These data suggest an important role in removing or repositioning Nuc3 in the activation of viral transcription by LRAs. The lack of increase in accessibility for the first two LTR peaks after LRA stimulation is in contrast with the behavior of the viral LTR in latently infected primary cells (Jefferys et al., 2021). This likely reflects the high baseline accessibility of the viral promoter in 2D10 cells and could contribute to the relatively high responsiveness of this cell line to LRAs.

We also examined the overall accessibility across the viral genome for different transcriptomic clusters, using the clusters assigned by graph-based clustering. Notably, cluster 6, which exhibited the highest level of viral transcription, showed the highest average accessibility across the HIV genome (**Figure 2F**). This observation is consistent with the hypothesis that an overall increase in accessibility across the HIV genome is correlated with increased viral gene expression, likely due to nucleosomal remodeling and histone modification to permit passage of RNA polymerase II through the provirus. Curiously, cluster 3 displayed the higher percentage of cells with HIV accessibility, but the average HIV accessibility was lower than for cluster 6.

### LRAs promote defined changes to the transcriptome and epigenome of HIV infected cells

To understand the impact of individual LRAs on host cells, we examined the host cell transcripts that were upregulated or downregulated by the addition of the LRAs. As expected, each LRA displayed a unique signature when considering the upregulated genes, with only three genes/transcripts exhibiting increased expression with all three LRAs (LEF1, FRYL, and HIV-LTR) (**Figure 3A**). The top differentially expressed genes are visualized in **Figure 3B**, and the full analysis results are included in **Table S1**. Prostratin had the largest impact on the host cell transcriptome (254 genes upregulated), consistent with its known ability to activate multiple signaling pathways, while vorinostat increased expression for 114 genes. By contrast, iBET151 had a more modest impact, with 56 genes being upregulated. Interestingly, the downregulated gene set exhibited significantly more overlap between the LRAs, with 134 genes showing decreased expression with all three LRAs. Notably, this core set of downregulated genes included many ribosomal proteins, suggesting a shared impact on cellular protein synthesis. To probe the biological significance of the differentially expressed genes, we performed Gene Ontology (GO) enrichment analysis to identify enrichment of specific functional classes of genes (**Table S2**). For the downregulated genes, we detected a shared set of enriched biological process terms across all three LRAs. Notably, one of the most highly enriched sets was viral transcription (GO:0019083), for which 64 genes were differentially expressed. These genes included mostly L ribosomal proteins (RPL) and S ribosomal proteins (RPS) genes, as well as UBA52 and FAU, confirming the regulation of ribosomal proteins on viral infection (Li, 2019). Interestingly, genes that are upregulated upon prostratin treatment were enriched in GO terms, including T cell receptor signaling pathway, as well as several pathways involved in signal transduction and regulation of cell migration (**Table S2**).

**Figure 3.**
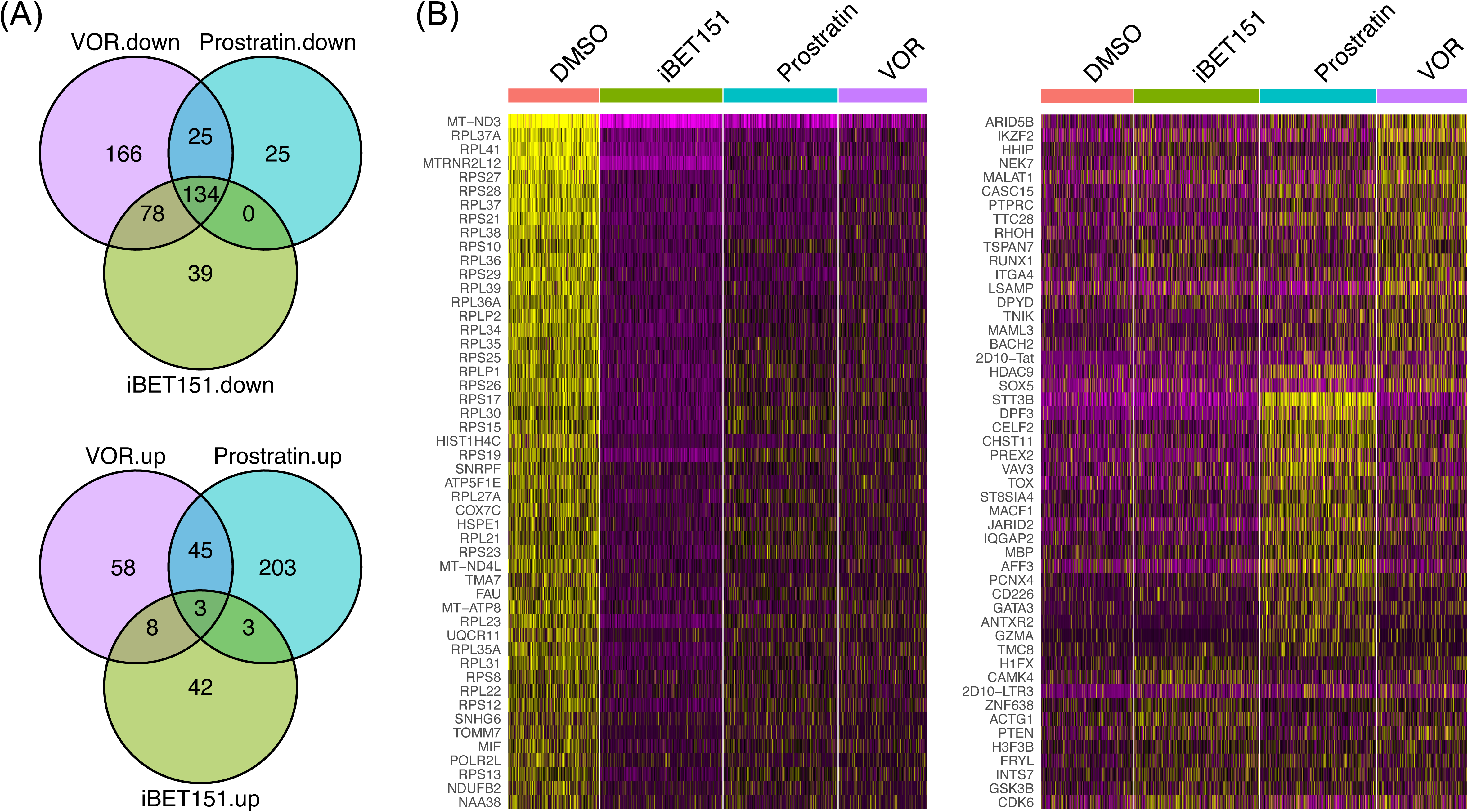
Differentially expressed genes by LRAs. **(A)** Venn diagrams showing the number of genes that were down-regulated (the top panel) and upregulated (the bottom panel) upon treatment with LRAs (FDR adjusted *p*-value ≤ 0.05). **(B)** Heatmaps showing normalized and scaled RNA expression levels for different treatment conditions. The rows contain representative genes that were down-regulated (left) and those that were upregulated (right), respectively. The columns represent single cells.

**Table 1.** Top 20 genes/TFs/peaks/enriched motifs linked with HIV viral expression. **(A)** Shown is a list of Spearman correlation coefficients (cor), nominal p-values from linear regression, and adjusted p-values (pval.adj) for top-ranked genes that were associated with HIV RNA expression. **(B)** Same as (A) but for TF linkage analysis. **(C)** Same as (A) but for peak linkage analysis. **(D)** A list of motifs enriched in ATAC-peaks that were significantly associated with HIV RNA expression. Corresponding TFs (TF), the number motif occurrences in the linked peaks and background peaks (observed, background), the percentage of linked peaks and background peaks containing the motif (percent observed, percent background), fold enrichment, and p-values from hypergeometric tests examining the degree of enrichment are shown as columns.

We next examined the impact of the LRAs on the cellular chromatin of HIV infected cells by using the scATAC-seq data to examine enrichment of TF binding sites within the open chromatin. During activation of a TF in a cell, TF binding to promoter and enhancer regions is frequently associated with an increase in accessibility at binding sites due to the action of nucleosomal remodeling complexes recruited by the TFs. Thus, increased enrichment of bindings sites for a TF in the open chromatin of a cell can serve as a proxy for overall activity of a TF in that cell. Using chromVAR (Schep et al., 2017) we calculated a TF deviation score for a set of 633 TFs with annotated binding motifs for each cell and examined the impact of each LRA on these TF scores via a test of differential motif accessibility. Similar to the scRNA-seq data, each LRA caused a distinctive impact on the pattern of TF activity in the host cells (**Figure 4A**). iBET151 had only a mild impact on the host cell TF scores, with 15 TFs increased and 15 TFs decreased, in concordance with the differential gene expression analysis. By contrast, vorinostat and prostratin increased the activity scores for 68 and 89 TFs, respectively. Notably, TF scores were significantly increased after prostratin stimulation for two members of the AP-1 family (JUN and FOS), consistent with this family’s known role in the PKC signaling pathway (**Figure 4B**). No TFs were increased by all three LRAs, but four TFs exhibited downregulated activity in response to all three LRAs; these TFs were FIGLA – a bHLH TF and three members of the SNAIL family of transcriptional repressors (SNAI1, SNAI2, and SNAI3) (**Figure 4C**).

**Figure 4.**
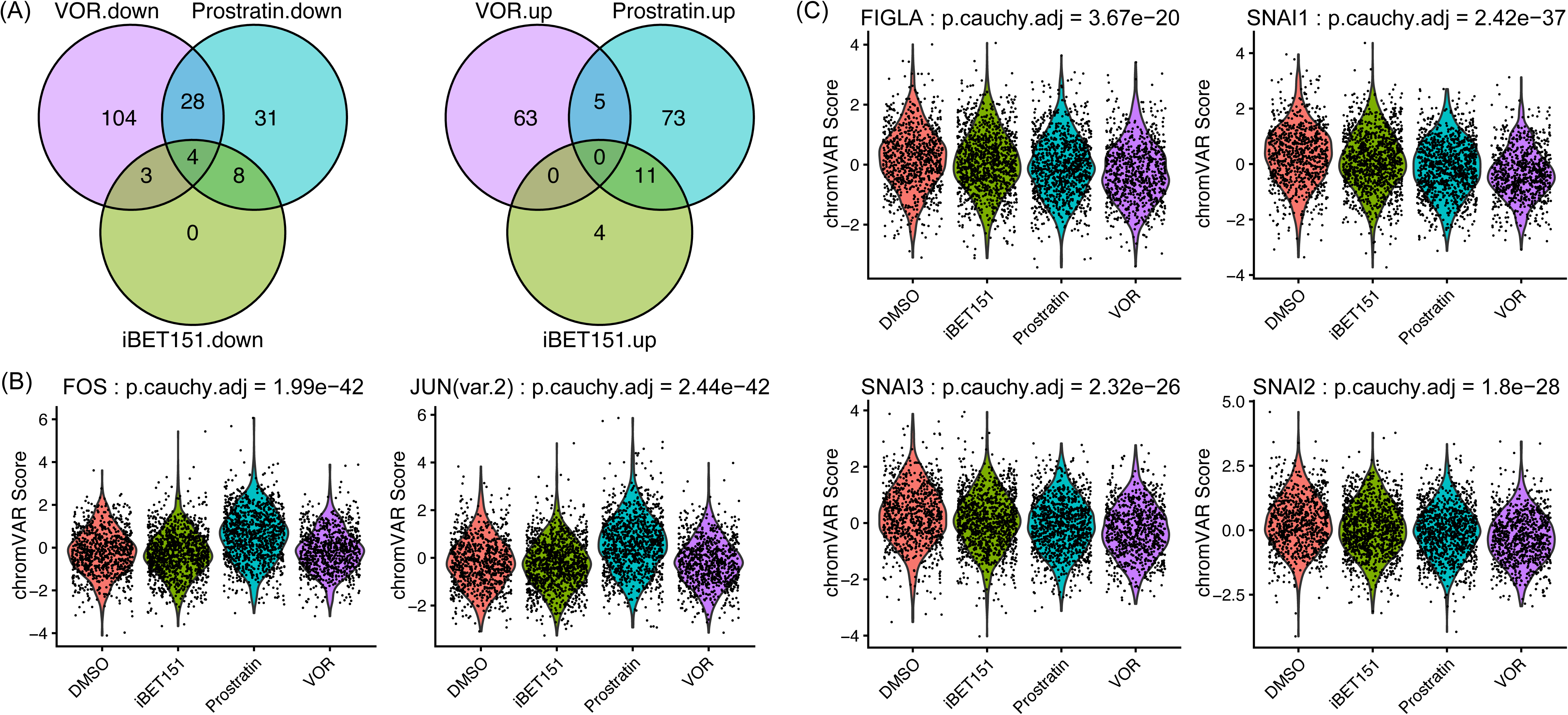
Differentially accessible TFs following LRA stimulation. **(A)** Venn diagrams showing the number of TFs whose motif accessibilities were significantly increased (the left panel) or decreased (the right panel) upon treatment with LRAs compared to DMSO (FDR adjusted *p*-value ≤ 0.05). **(B)** Violin plots showing distributions of motif accessibilities (deviation scores) of positive controls, FOS and JUN. *p*-values from Wilcoxon rank sum tests comparing DMSO and one of the LRA-treated samples were integrated using the Cauchy combination. **(C)** Same as (B) but for the four intersected TFs from (A): FIGLA, SNAI1, SNAI3, and SNAI2.

### Identification of cellular transcripts that correlate with HIV RNA levels

To understand the relationship between vRNA, cellular RNA, and chromatin accessibility more deeply, we examined pairwise correlations between summed vRNA levels for each cell and individual cellular transcript levels across cells of the dataset. We performed this analysis for each individual condition, as well as across the aggregated cell population. Although we found some condition-specific associations, we observed that including all conditions in this analysis identified a significantly greater number of HIV-correlated transcripts, likely due to the increased cellular diversity and statistical power provided by the larger cell numbers (**Figure S7, Figure S8**). Therefore, we focused our analysis on the aggregated cell population. We identified 229 and 123 cellular transcripts whose expression correlated positively or negatively with HIV vRNA levels using a cutoff of FDR-adjusted *p*-value ≤ 0.05, respectively (**Figure 5A, 5B**). The top significantly linked genes are included in **Table 1A**, with the full testing results shown in **Table S3**. Overall, each transcript in this set individually correlated with HIV vRNA levels only weakly, albeit the small *p*-values, with a 95% confidence interval of [0.001, 0.0159] for the significant *R*^2^. As expected, the most highly correlated transcripts were individual viral RNAs. The most highly correlated cellular transcript was TSPOAP1, a cytoplasmic protein with no previously known role in HIV replication. Although the cellular role of TSPOAP1 is unclear, it has recently been shown that TSPOAP1 enhances influenza A virus replication by antagonizing innate immune signaling (Wang et al., 2022). Interestingly, two of the most strongly correlated transcripts were long noncoding RNAs (MALAT and NEAT) that have both been previously implicated in HIV replication (Qu et al., 2019). In particular, MALAT1 has been shown to negatively regulate the PRC2 complex responsible for adding repressive H3K27me marks to histones. This mark has been shown to be associated with repressed HIV transcription, and PRC2 inhibition promotes HIV gene expression (Friedman et al., 2011; Turner et al., 2020). Another lncRNA (ZFPM2-AS1) with no known role in HIV was also positively correlated with HIV RNA levels. The list of positively correlated transcripts also included many genes with no previous known role in HIV gene expression, including ANXA1 (Galvao et al., 2020), which regulates innate immune signaling, and CELF2, which regulates T cell development and activation (Mallory et al., 2015).

**Figure 5.**
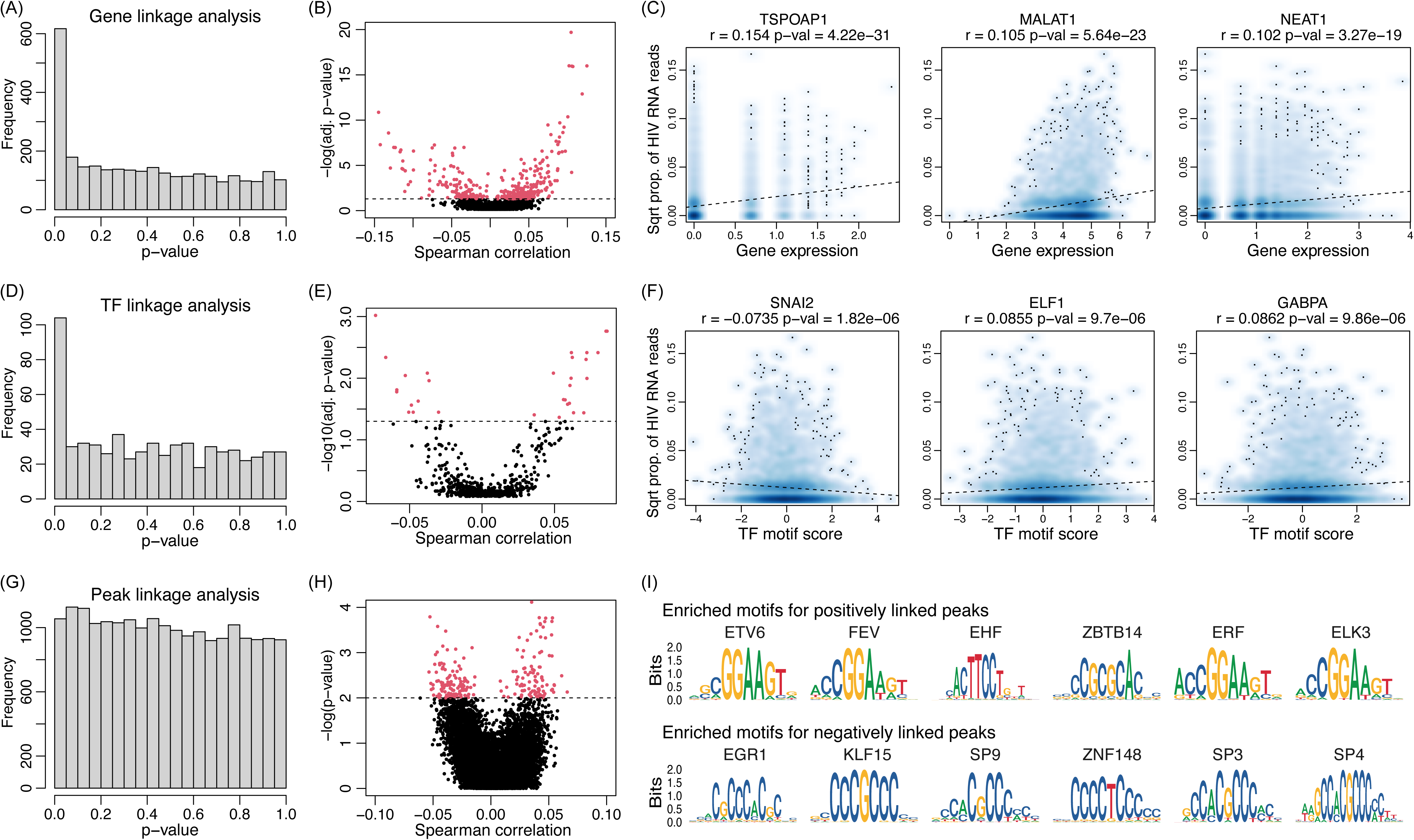
Linkage analysis between HIV viral expression and genes/TFs/peaks. **(A)** Distributions of nominal *p*-values from gene expression linkage analysis. **(B)** Volcano plot showing the negative log transformed adjusted *p*-values against the Spearman correlation coefficients from gene expression linkage analysis. The dotted line represents the adjusted *p*-value of 0.05. Points with *p*-values below this threshold are colored in red. **(C)** Visualizations of top linked genes. The dotted line represents the fitted line from a simple regression; Spearman correlation coefficient and nominal *p*-value are included in the title. **(D)** Same as (A) but for TF activity linkage analysis. **(E)** Same as (B) but for TF activity linkage analysis. **(F)** Same as (C) but for TF activity linkage analysis. **(G)** Same as (A) but for peak linkage analysis. **(H)** Same as (B) but for peak linkage analysis. Due to the large number of testing and the ATAC sparsity, we used the nominal *p*-values instead. **(I)** Visualization of motifs enriched in ATAC peaks that were positively (top) and negatively (bottom) associated with the HIV RNA expression.

Notably, many of the cellular transcripts that correlated negatively with HIV gene expression encoded ribosomal proteins. For example, the transcript most negatively correlated with HIV was RPL36, followed by RPL37A and RPS27 (**Table S3**). GO enrichment analysis using the negatively linked genes identified enriched terms, including viral transcription and regulation of cell cycle (**Table S4**). Indeed, transcription of ribosomal proteins is typically variable depending on the phase of the cell cycle (Nosrati et al., 2014). Therefore, we further examined whether HIV transcript levels were associated with cell-cycle marker genes, using known signatures for the S phase and G2-M phase to calculate a phase “score” for each cell, along with the predicted classification of each cell in either G2M, S or G1 phase. We observed a significant association between HIV RNA expression and cell-cycle classifications (one-way ANOVA *p*-value = 6.6e-4) (**Figure S9A**). We observed a modest negative correlation between HIV gene expression and expression of S phase genes (r = −0.071, *p*-value = 1.027e-5; **Figure S9B**), and no correlation with G2-M expression (**Figure S9C**). Overall, this suggests that cell cycle phenomena may make a modest contribution to the induction of HIV by LRAs.

RNA levels for several transcription factors were found to be correlated with HIV RNA (**Figure S10**). GATA3 was the most significantly positively correlated TF (*p*-value = 0.0005), and interestingly, the chromatin insulator CTCF was the most significant negatively associated factor (*p*-value = 0.0072). We have recently reported that CTCF is a novel repressor of HIV gene expression in CD4 T cells, and knockdown of CTCF in latently infected Jurkat cells and primary CD4 T cells reactivates HIV gene expression (Jefferys et al., 2021). YY1 and HSF1, two TFs that have been confirmed to repress HIV replication, were also found to be negatively correlated with HIV RNA levels (Coull et al., 2000; Nekongo et al., 2020). Overall, these data indicate that this approach is able to successfully identify known regulators of HIV expression and reactivation, but also reveals novel HIV associations with cellular transcripts that could represent important but uncharacterized viral regulators.

### Activity of individual TFs correlates with HIV gene expression

The integrated multiomic approach also allowed us to examine connections between the epigenomic state of the cells and viral gene expression. Using the chromVAR motif scores calculated from the scATAC-seq reads for each cell, we also investigated whether activity of individual TFs correlated with HIV RNA levels across the cell population. We expect that TFs that promote HIV gene expression will have motif deviation scores that positively correlate with HIV RNA levels, while TFs that repress HIV will have deviation scores that negatively correlate with HIV RNA. We observed that 31 TFs had motif scores that are positively or negatively correlated with HIV RNA using a cutoff of FDR-adjusted *p*-value ≤ 0.05 (**Figure 5D, 5E, Table 1B**). Similar to the correlations between HIV RNA and the abundance of cellular transcripts, the overall correlation coefficients between HIV RNA and TF activity scores, although statistically significant, were individually weak (95% confidence interval of [0.0012, 0.0073] for the *R*^2^). Condition-specific analyses, albeit underpowered, could also recapitulate the associations from the aggregated analysis (**Figure S11**).

Among TFs positively correlated with HIV expression, multiple members of the ETS, GATA and AP-1 families were identified. Notably, the HIV LTR promoter contains binding sites for members of all three of these families, and previous work has shown that individual members of these families can positively regulate HIV gene expression, indicating that this approach successfully identifies HIV regulators. For example, the HIV LTR promoter contains two ETS binding sites and the ETS TFs ETS-1, ETS2, ELF1 and GABPA have been previously identified as regulating HIV gene expression (Flory et al., 1996; Hilfinger et al., 1993; Panagoulias et al., 2017; Yang et al., 2009). Similarly, AP-1 has at least four binding sites in the LTR and this TF family is potently activated by PKC agonists (Rabbi et al., 1997). Furthermore, our lab has previously determined that elevated activity of GATA TFs was associated with increased viral gene expression (Jefferys et al., 2021). However, the role of specific members of the ETS, GATA and AP-1 families from these data should be interpreted with some caution, given that these members bind to highly similar sequences and accurately distinguishing the activity of them can be challenging.

Amongst the negatively correlated TFs, two families of TFs were prominently represented. In particular, three members of the TCF family were associated with lower HIV expression – TCF3, TCF4 and TCF12. Some previous data has linked a related TCF protein (TCF1) to HIV regulation through direct binding (Waterman and Jones, 1990). Additionally, TCF4 has been shown to repress HIV expression (Henderson et al., 2012; Wortman et al., 2002). Interestingly, our analysis also identified members of the SNAIL family of transcriptional repressors as being negatively correlated with HIV gene expression (SNAI1, SNAI2 and SNAI3). This family of TFs binds to E-BOX motifs and have not been previously implicated in HIV expression. Notably, four potential E-BOX motifs (CANNTG) are present in the HIV LTR. Other TFs with no known role in HIV regulation were also identified −TBX5 and TBX15 are T box transcription factors that bind T box motifs (TCACACCT), and two potential T box motifs are present in the HIV genome (nts 5433-4442 and 8995-9002).

We also examined the correlation between accessibility at individual cellular chromatin peaks and viral transcription. In general, correlation coefficients between individual peaks were even weaker than for transcripts and for TF scores, with typical spearman correlations between 0.02 and 0.05 (**Figure 5G, 5H, Table 1C**). Note that the statistical power is low due to the large number of testing/peaks and the data sparsity (**Table S5**). Nevertheless, we detected 224 peaks whose accessibility was significantly correlated with HIV transcript levels, with a significance threshold of 0.01 on the nominal *p*-values. We then examined this set of positively and negatively linked peaks for motif enrichment compared to a set of background peaks using Signac. We found significant enrichment of motifs that represented binding sites for 91 TFs (**Table 1D**). Members of the Sp and Kruppel-like factors (KLFs) were prominently represented in this list Family included Sp4, Sp1 and Sp9, as well as KLF15, EGR1, and ZNF148 (**Figure 5I**).

We then examined overlap between sets of TFs that were detected as significantly associated with HIV RNA levels across multiple analyses (linked expression, linked TF score and enrichment in linked peaks). We found ten TFs that were detected as significantly associated with HIV across all three sets (**Figure 6A; Table S6**). Additionally, to examine coordinate behavior between HIV associated TFs, we examined pairwise correlations of consensus principal components derived from both TF expression and deviations scores of TF binding motifs, within a set of top 50 TFs that have the smallest *p*-values from the Cauchy combination test (**Table S7**). An adjacency matrix was constructed for pairs with correlation coefficients of > 0.5 (**Figure 6B**). The data indicated that this set of TFs could be grouped into three distinct clusters (**Figure 6C**). One cluster was composed mainly of KLF and Sp TFs but also included chromatin insulators CTCF and CTCFL. The second cluster was composed almost entirely of ETS TFs, while a third cluster contains just three TFs that are members of the SNAIL family of transcriptional repressors – SNAI1, SNAI2 and SNAI3. While the KLF/Sp TFs in general behaves as a single tightly connected unit, some substructure was evident in the ETS cluster, with 3-4 groups forming more tightly connected subclusters within the overall cluster. Overall, these results from TF network analysis shed light upon the regulatory relationships and synergies among the previously identified TFs.

**Figure 6.**
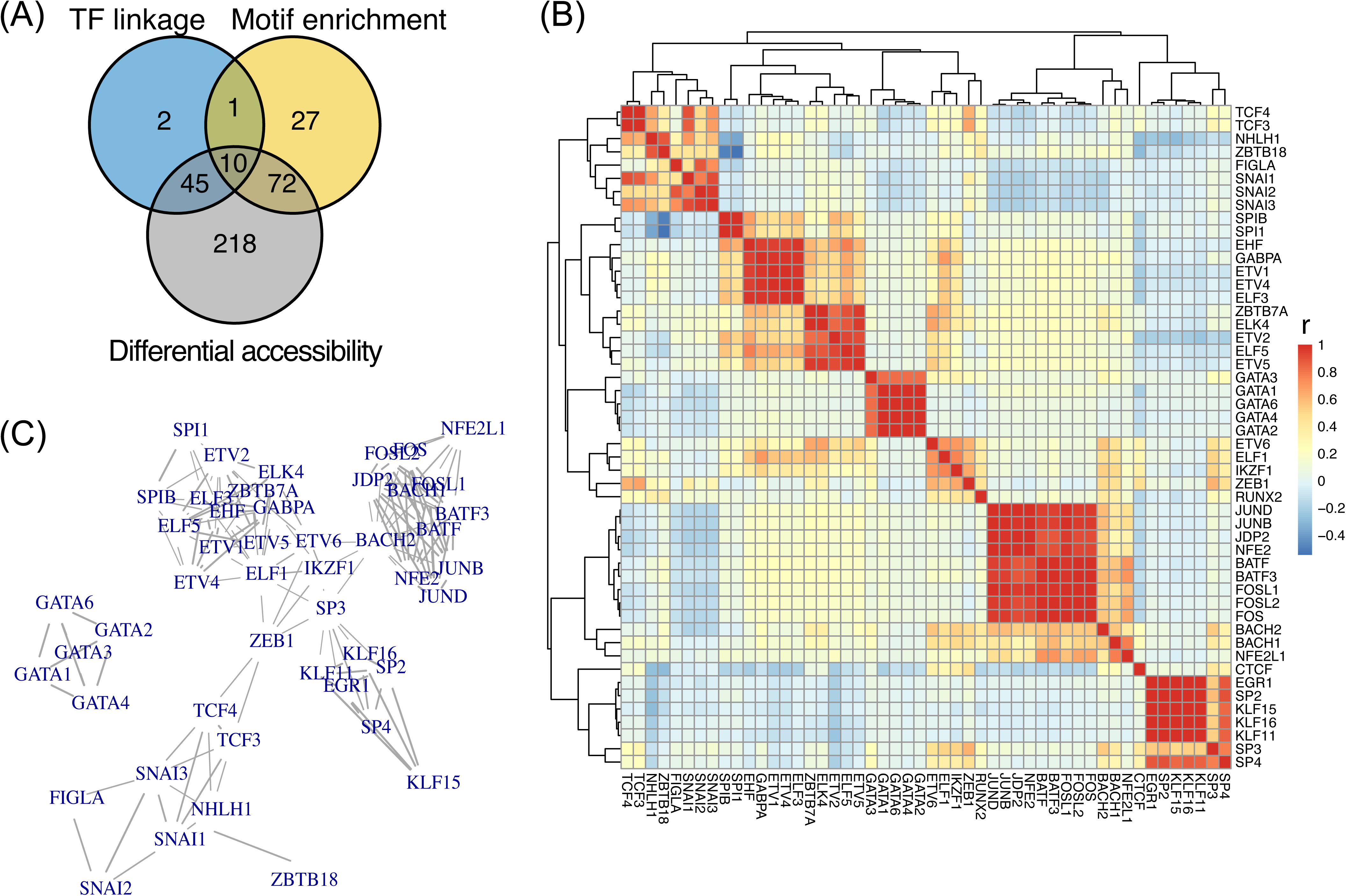
TF regulators of HIV latency reversal with experimental validations. **(A)** A Venn diagram showing the number of significant TFs from three different testing schemes: TF linkage analysis, enrichment analysis of TF-binding motifs using significantly linked peaks, and analysis of differential motif accessibility. **(B)** Heatmap showing correlations between the TF pairs. The correlation is computed using the TF-specific consensus principal components, calculated using both the TF expression and its motif accessibility. **(C)** TF regulatory network.

### Machine learning model of HIV transcription from multiomic data

Since each transcript, TF score, or peak only weakly correlated with HIV RNA levels, we next examined whether considering multiple features of the dataset at once in a multivariate machine learning model could lead to a more predictive model of HIV gene expression from the dataset (**Figure 7**). We first used Spearman correlation coefficients between individual features (transcripts, TF scores, peaks) and HIV RNA levels to select the 60 highest ranked features and then incorporated these features into a boosting-based machine learning model. To simplify the prediction goal of the model, we converted the data from the cells into a binary classification of HIV RNA+ and HIV RNA-based on whether HIV RNA was detected in each cell. For machine learning training and evaluation, we split the data into five sets – four for separate training and one for testing. By examining the receiver operating characteristic (ROC) curves to evaluate model performance, the model was able to achieve an area under the curve (AUC) of 0.79 for the training set and 0.68 for the test set (**Figure 7A**). We were also able to determine the rank importance of individual features to the model performance (**Figure 7B**). Based on this ranking, the most important predictive features were the transcripts TSPOAP1 and MALAT1 followed by accessibility of binding sites for the ETS TF SPIB (MA0081.2). The binding motif of another ETS TF (ELF1) was also ranked on the top ten (MA0473.3). Only one individual peak was present within the top 10 most predictive features - localized to H4C4 gene. Interestingly, transcript levels for the gene MSRB1 contributed this the performance of this model. The MSRB1 gene represents the integration site for the provirus in this cell line (Friedman et al., 2011; Pearson et al., 2008), suggesting a relationship between transcription of the provirus containing gene and viral reactivation.

**Figure 7.**
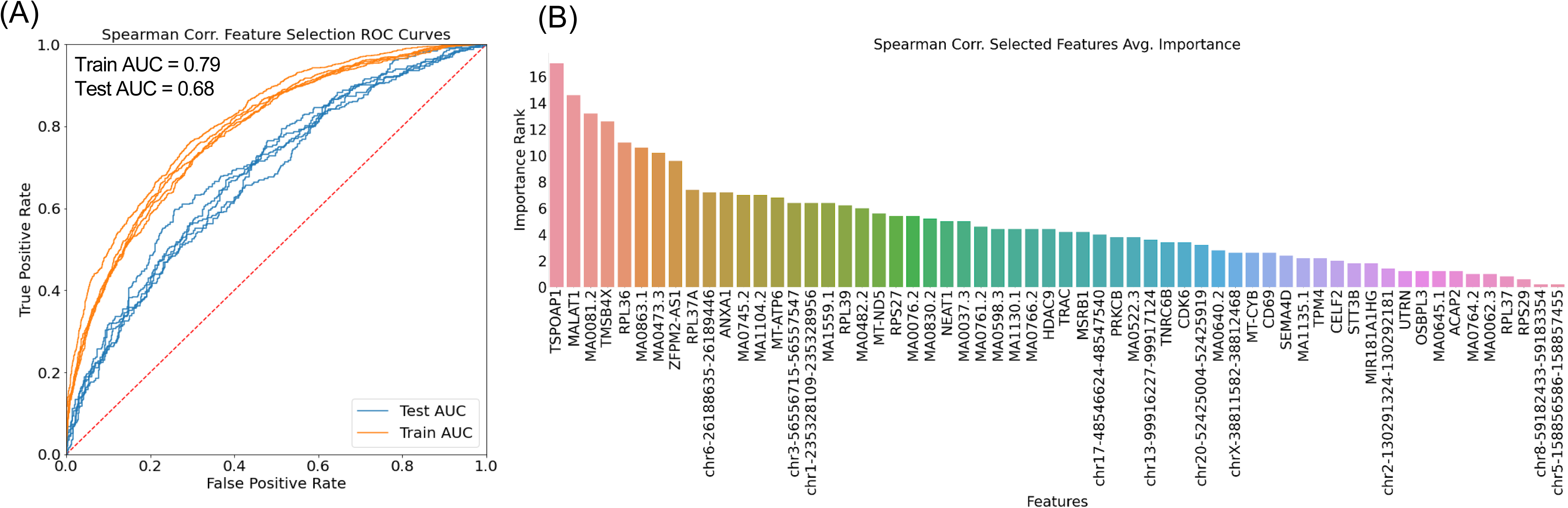
Machine learning model of HIV reactivation. **(A)** Receiver operating characteristic (ROC) curve for a machine learning model of HIV expression using the single cell multiomic dataset. The model was trained on a subset of the overall data, then tested on the remainder. Representative curves from the training (orange) and test (blue) sets are shown. Area under the curve (AUC) is labeled. **(B)** Ranking of model features based on overall contribution to model performance.

### GATA3 regulates HIV latency

Based on these analyses, we chose to further investigate the role of the transcription factor GATA3 in HIV latency. Our scRNA-seq data had identified the transcript for GATA3 as being positively correlated with HIV vRNA levels and the scATAC-seq data indicated that activity of GATA3 was positively correlated with HIV vRNA levels. Furthermore, GATA3 motif (MA0037.3) accessibility played a significant role in the performance of our machine learning model of HIV reactivation (**Figure 7B**). Thus, we hypothesized that GATA3 activity positively regulates HIV gene expression and that it plays an important role in the response to one or more of the LRAs in this study. We first examined the activity and expression of GATA3 across the conditions. We observed that GATA3 expression and activity were both significantly increased over DMSO control in the prostratin stimulated cells and, to a lesser extent, in VOR treated cells (**Figure 8A, 8B**). We further investigated the role of GATA3 in HIV latency by transducing 2D10 cells with a GATA3-targeting shRNA lentivirus or a control shRNA lentivirus. Transduced cells were selected by puromycin, and efficient knockdown of GATA3 was confirmed by western blot (**Figure 8C**). The cells were then stimulated with 500nM prostratin to reactivate HIV and viral gene expression measured by flow cytometry for GFP. We observed that cells transduced with the GATA3-targeting shRNA exhibited a significantly reduced response to prostratin (~44%) compared to untransduced cells or cells with a non-targeting shRNA (77% and 67%) respectively (**Figure 8D**). Although GATA3-independent pathways clearly account for a significant fraction of the response to prostratin, these data demonstrate that GATA3 plays an important role in viral reactivation by this molecule. Overall these findings highlight the value of the combined multiomics approach to identify and prioritize candidate new regulators of HIV and reveal GATA3 as a previously unappreciated factor that impacts the reactivation of HIV in response to LRAs.

**Figure 8.**
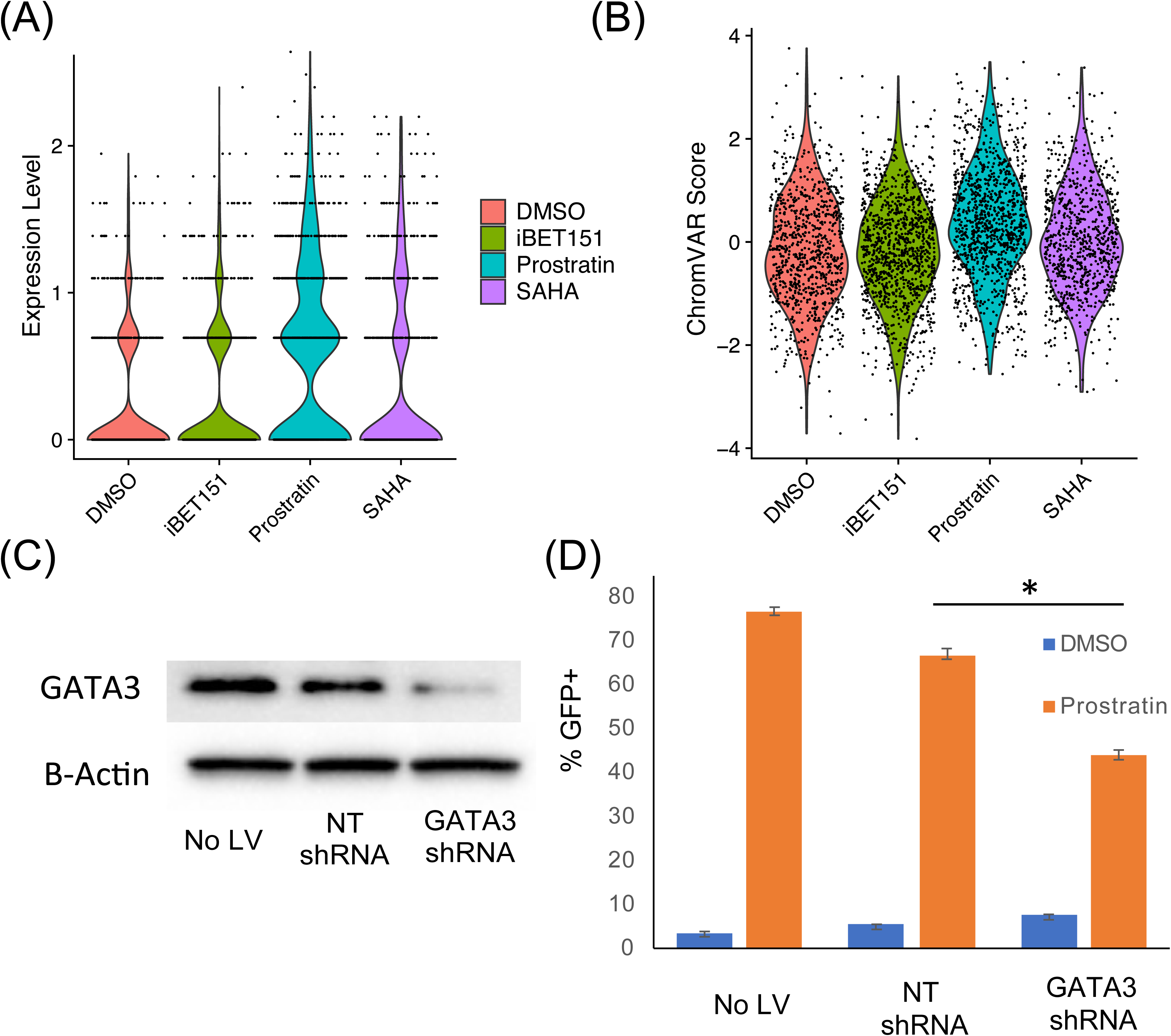
GATA3 regulates HIV latency. **(A)** Violin plot of GATA3 expression across experimental conditions, data taken from scRNA-seq data. **(B)** Violin plot of GATA3 TF activity across experimental conditions based on chromVar analysis of scATAC-seq data. **(C)** 2D10 cells were transduced with a GATA3-targeting shRNA or a control non-targeting shRNA (NT shRNA), selected with puromycin, then protein extracts analyzed by western blot for Beta-Actin and GATA3. **(D)** shRNA transduced 2D10 cells were stimulated with 500nM prostratin for 24h. Viral reactivation was measured by flow cytometry for GFP expression. Data represent the average of triplicate readings. Asterisks indicate P<0.05, Students T Test.

## Methods

### Reactivation of latently infected 2D10 cell line

A latently infected Jurkat derived T-cell line (2D10 cells) was cultured in RPMI 1640 supplemented with 10% FBS, penicillin (100 IU/ml), streptomycin (100 μg/ml), and 25 mM HEPES at 37°C in 5% CO2. The 2D10 cell line carries an HIV provirus that expresses the regulatory proteins Tat and Rev, and a short-lived green fluorescent protein (d2EGFP) in place of Nef. The latently infected cells were reactivated with three latency-reversing agents (LRAs) with different mechanisms of action, prostratin (PKC activator), iBET151 (Bromodomain inhibitor), or vorinostat (HDAC inhibitor). Concentrations used for reactivation of the virus were 75nM for prostratin and iBET151 and 500nM for vorinostat. 24h post treatment, reactivated 2D10 cells were quantified for GFP expression by flow cytometry using a Becton Dickson Fortessa and analysis using FlowJo software.

### Single-cell multiome (scRNA-seq and scATAC-seq) library construction and sequencing

Construction of single cell multiome libraries was performed using a 10XGenomics Chromium controller and a single cell multiome ATAC + Gene expression kit following manufacturers protocol. Nuclei were first isolated following the 10XGenomics protocol (CG000365 • Rev B). Then, nuclei suspensions were subjected to Tn5 transposition in bulk, followed by barcoding using GEM (Gel Beads-in-emulsion) beads. Silane magnetic beads were used to purify the barcoded products from the post GEM-RT reaction mixture. Incubation of the GEMs produces barcoded DNA from the transposed DNA (for scATAC-seq) and 10x Barcoded, full-length cDNA from poly-adenylated mRNA (for scRNA-seq). Barcoded transposed DNA and barcoded full-length cDNA were then pre-amplified by polymerase chain reaction (PCR) to facilitate library construction. The pre-amplified product was used as input for both ATAC and gene expression (GEX) library construction. P5 and P7 indexes were added to the pre-amplified transposed DNA for ATAC-seq library. cDNA amplification, enzymatic fragmentation followed by end Repair, A-tailing, adaptor Ligation, and PCR were performed to incorporate P5, P7, i7 and i5 sample indices, and TruSeq Read 2 (read 2 primer sequence) for gene expression libraries. The libraries were quantified using an Agilent Tapestation 4200 and the Qubit dsDNA High Sensitivity Assay Kit (Invitrogen, #Q33230). Pooled samples from each ATAC and RNA libraries were sequenced using paired-end, single-index (ATAC-seq) and dual index (RNA-seq) sequencing on a NextSeq 2000 instrument (Illumina). For GEX libraries, the read format was: Read 1 - 28 cycles; Read 2 - 90 cycles; i7 - 10 cycles; i5 – 10 cycles. For ATAC libraries the read format was: Read 1 – 50 cycles; Read 2 – 49 cycles; i7 – 8 cycles, i5 – 16 cycles. Paired end reads of pooled libraries were demultiplexed prior to downstream analysis.

### scRNA-seq and scATAC-seq data processing

For the 10XGenomics Multiome data, we used Cell Ranger ARC (version 2.0.0) to perform sample demultiplexing, barcode processing, peak calling, and counting of RNA and ATAC reads in single cells. The HIV genome is included as a separate chromosome, with HIV genes and peaks shown in the coverage plots in **Figure 2E** and **Figure S4**. For scRNA-seq and scATAC-seq, 36,607 genes and 71,335 peaks were profiled across 5,333 cells (1,124 DMSO, 1,455 Prostratin, 1,186 VOR, and 1,568 iBET151), respectively. After quality control procedures on total ATAC reads, total RNA reads, and mitochondrial percentages, 3,900 cells (850 DMSO, 1,071 Prostratin, 829 VOR, and 1,150 iBET151 cells) were kept for further analyses. For read count normalization, we used sctransform by Seurat (Stuart et al., 2019) for scRNA-seq and TF-IDF by Signac (Stuart et al., 2021) for scATAC-seq, respectively. This is followed by principal component analysis (PCA) for dimension reduction and UMAP (Becht et al., 2018) for visualization. For scATAC-seq data, we also adopted weighted PCA by Destin (Urrutia et al., 2019) weighing chromatin accessibilities by existing genomic annotations and publicly available regulomic data. We further adopted the weighted nearest neighbor method by Seurat (Hao et al., 2021) to jointly identify cell clusters across both modalities (**Figure 1C**).

We obtained the position frequency matrices and annotated 633 pairs of TFs and motifs from the JASPAR database (Fornes et al., 2020); we further applied chromVAR (Schep et al., 2017) to derive, for each TF, its motif score, which measures the deviation in chromatin accessibility across the set of peaks containing the corresponding motif, compared to a set of background peaks. We also constructed a pseudo-gene activity matrix (Pliner et al., 2018), summing ATAC read counts in gene bodies and promoter regions (2Kb upstream of genes’ transcription start sites), followed by sctransform normalization.

### Differential gene expression and motif accessibility

For each treated group (iBET151, Prostratin, and VOR), we performed a nonparametric Wilcoxon rank sum test to identify differentially expressed genes (**Figure 3A**) and differentially accessible motifs (**Figure 4A**) compared to the DMSO control. We used the normalized gene expressions by sctransform and the motif deviation scores by chromVAR as input, and adopted FDR for multiple testing correction. Cauchy combination testing (Liu and Xie, 2020) was performed to integrate the three sets of condition-specific *p*-values that are correlated (**Figure 4B, 4C**). For significantly differentially expressed genes, we carried out a gene ontology (GO) enrichment analysis using DAVID (Huang da et al., 2009).

### Gene, TF, and peak linkage analysis

For HIV expression, we calculated, for each cell, the proportion of HIV RNA reads out of the total number of reads. For HIV chromatin accessibility, we directly used the total number of HIV ATAC reads, due to the sparsity of the scATAC-seq data. The correlation between the HIV expression and HIV accessibility was marginally significant (**Figure 2G**). We used the squared root of the proportion of HIV RNA reads as the outcome variable in various supervised frameworks, aiming to recover the molecular biomarkers that are related to HIV expression upon LRA treatment and thus latency reversal. The dependent variables in such linkage analyses include gene expression (**Figure 5A, 5B**), TF motif scores (**Figure 5D, 5E**), and peak accessibilities (**Figure 5G, 5H**). For gene and TF linkage analyses, we adopted FDR to control for false positives and to adjust *p*-values. Due to the large number of testing in the peak linkage analysis and the low power due to the ATAC sparsity, we used a significance threshold of 0.01. For the significantly linked peaks, we further searched for motifs that are overrepresented in them compared to a background peak set using the hypergeometric test of enrichment. We repeated the linkage analyses in a condition-specific manner, and the results, albeit with lower power due to the lower number of cells in each treated group, are generally reproducible (**Figure S7**). To examine associations between HIV expression and cell cycle, we computed cell cycle scores as implemented by Seurat (Stuart et al., 2019). The HIV genes were excluded when a control set was selected.

### TF network analysis

For TF-specific testing from the TF linkage analysis, motif enrichment analysis, and differential accessibility analysis, we compared the calling results (**Figure 6A**) and generated a master output to identify key TF regulators associated with latency reversal (**Table S7**). We performed consensus PCA to jointly reduce dimensions of TF expression and deviation scores of TF binding motifs. For each pair of TFs, we calculated the correlation coefficients between their consensus principal components (**Figure 6B**) and constructed an adjacency matrix for the TF network by creating an edge between two TFs if the correlation between them is greater than 0.5 (**Figure 6C**).

### Machine learning model of HIV reactivation

Due to the large number of features in the dataset, feature selection was necessary to create an interpretable and usable set of features for a machine learning model. In order to select a high-performing subset of features for prediction, we first calculated Spearman correlation between each feature within the multiomic dataset against the HIV gene expression before binarization of the cell data to HIV+ or HIV-based on the presence or absence of any detectable viral RNA. From each feature set (RNA expression, motifs, and peaks), features with top k ranking Spearman correlation scores were then selected for modeling. Here k is a hyperparameter to be determined by parameter tuning. After careful tuning, we chose 30 features from RNA expression, 20 from motifs and 10 from peaks. These selected features were then used to train a gradient-boosting tree classifier, which constructs a strong final classifier by combining several weaker classifiers. In gradient boosting, weak learners are optimized in an iterative fashion with loss functions that focus on the residuals (errors) of their predecessors. Here we used LightGBM, an efficient implementation of this algorithm (Ke et al., 2017). The parameters of this model were then tuned via 10-fold cross-validation and grid search. The feature importance in LightGBM is calculated as the total number of splits of each feature in all trees and was used in our experiment to rank each selected input feature.

### GATA3 shRNA knockdown

An shRNA lentivirus targeting GATA3 was obtained from The RNAi Consortium (TRCN0000019303) along with a non-targeting parental lentiviral vector (pLKO). For the shRNA lentiviruses, backbone plasmids were cotransfected into HEK293T cells with GagPol and VSV-G packaging plasmids to generate lentiviral particles. Virus containing supernatant was subjected to low-speed centrifugation (300g, 5min) and filtration using a 0.45um filter to removed cellular contaminants. The remaining supernatant was added to media with 2D10 cells for 24h to facilitate infection. For shRNA transduced cells, we selected for transduced cells by adding 1ug/mL puromycin to the tissue culture media for 4-5 days. Western blotting for GATA3 and Beta-Actin was carried out by lysing the cells with RIPA buffer (ThermoFisher) supplemented with benzonase and protease inhibitors. Protein extracts were quantified by Bradford assay (BioRad) and run on a 4-20% Tris-Glycine polyacrylamide gel (ThermoFisher), before transfer to a nitrocellulose membrane. Primary antibodies for detection were anti-GATA3 (abcam) and anti-beta actin (abcam). Washing was carried out using Tris-buffered saline, with 0.1% Tween-20 detergent. Targets were then detected using horse-radish peroxidase linked secondary antibodies (Novex) and developed using ECL reagents (ThermoFisher) before imaging on a BioRad gel imager.

## Discussion

Inefficient reactivation of HIV by current latency reversing agents is a major impediment to achieving an HIV cure, and developing methods that can broadly reactivate the replication competent reservoir is an urgent priority (Ho et al., 2013). However, the mechanisms responsible for this inefficiency remain largely unknown. Reactivation is regulated in part by stochastic processes relating to fluctuations in the abundance of host cell factors and in the level of the viral trans-activator Tat (Razooky et al., 2015; Weinberger et al., 2005). Nevertheless, reactivation is also likely influenced by the overall host cell environment, through the expression or activity of cellular transcriptional regulators that bind and directly regulate the viral LTR, or that influence cell fate decisions that indirectly impact viral gene expression (Roebuck and Saifuddin, 1999). Previous work from our lab and others has demonstrated that viral silencing in CD4 T cells is not random but is associated with a distinct transcriptomic and epigenomic program (Bradley et al., 2018; Golumbeanu et al., 2018; Jefferys et al., 2021). We hypothesize that during the initiation of HIV latency, a set of heritable epigenetic modifications are made to the provirus that maintain viral silencing through cell division, and that the frequency with which these modifications occur is connect to the overall host cell program (Turner and Margolis, 2017). Furthermore, this program, in combination with stochastic responses to latency reversing drugs likely regulate the response of latent proviruses to external latency reversing stimuli (Dar et al., 2014). Rather than representing a single state, we propose that latency represents a set of adjacent and interconnected states each characterized by a different set of restrictions to full viral reactivation. The low efficiency of overall reactivation for a given LRA is thus explained by the fact that the pathway targeted by each LRA is limiting only a minor fraction of cells (responders) while for other cells (non-responders) a different set of restrictions continue to block viral expression. As such, detailed investigation of viral reactivation at the single cell level may reveal additional cellular factors that control reactivation and could be targeted in parallel with current LRAs to greatly improve the overall potency and breadth of LRAs.

Understanding the heterogeneous and dynamic nature of HIV latency will require the application of single cell methods to model systems of latency in which, ideally, multiple modalities regarding the state of the host cell and the provirus can be simultaneously measured (Rato et al., 2017). By combining high dimensional, multimodal observations of latency reversal with analytical methods to connect these observations to viral gene expression, we aim to reveal the complex relationship between host cell factors and viral reactivation. In particular, combining scRNA-seq with scATAC-seq will allow us to connect viral expression to the behavior of cellular chromatin, and, by inference, the activity of individual cellular transcription factors (Stuart et al., 2019). Since TFs frequently recruit chromatin remodeling complexes to cellular promoters and enhancers, the re-opening of cellular chromatin in response to stimulation can reveal activity levels for sequence specific cellular TFs (Buenrostro et al., 2015; Schep et al., 2017). This paper represents the first use of combined scRNA-seq/scATAC-seq to analyze HIV infected cells.

In this study, we reactivated an established cell line model of latently infected cells with three different latency reversing agents in parallel, using concentrations that produce variegated reactivation of the provirus. As expected, each LRA induced a distinct set of gene expression and TF activity across the population. The overall diversity of transcriptomic and epigenomic phenotypes across the cell population then allowed us to examine the correlation of proviral accessibility and expression with numerous host cell factors. The overall expression of HIV transcripts was significantly correlated with proviral accessibility, consistent with the hypothesis that accessibility of the provirus is a key barrier to reactivation of expression. Curiously, LRA stimulation did not cause a significant enhancement to accessibility of the viral LTR promoter region in the stimulated cells. This observation is in contrast with bulk ATAC-seq observations of LRA stimulated primary cells, in which significant increase in promoter accessibility was observed (Jefferys et al., 2021). Instead, we observed that in LRA treated 2D10 cells, a peak located downstream of the viral transcription start site (TSS) increased accessibility in LRA stimulated conditions. We speculate that the relatively ‘easy-to-reactivate’ phenotype of the 2D10 cell line may be due to high baseline accessibility at the viral promoter and that chromatin remodeling at the viral promoter does not represent a major barrier to HIV reactivation in 2D10 cells (Kim et al., 2011). By contrast, the major difference in proviral accessibility in cells with high levels of viral transcription represented increased accessibility across the gene body of HIV, indicating that increased opening across the provirus is important to viral transcription.

By examining the correlation of cellular transcripts and TF activity with vRNA, we were able to identify a set of 348 differentially expressed cellular transcripts and 31 differentially active cellular TFs that significantly correlated (P<0.05) with vRNA levels across the whole population. Furthermore, we were able to examine the correlation of accessibility of specific chromatin peaks with HIV vRNA and examine these for TF site motif enrichment. Altogether, this approach has identified a set of candidate regulators of HIV expression and reactivation that could be investigated further for functional roles in HIV infection. Reassuringly, this approach identified several known HIV regulators of HIV transcription and latency including AP-1, Sp1, CTCF, and MALAT1 (Mbonye and Karn, 2017; Siliciano and Greene, 2011; Van Lint et al., 2013). This observation suggests that this approach is powered to detect real regulators of HIV transcription. In addition to these known HIV regulating factors, many genes or TFs with no known connection to HIV transcription were found. These included genes/TFs that either were either positively correlated with HIV transcription (candidate activators) or negatively correlated (candidate repressors). Amongst the positively correlated transcription factors, ETS and GATA transcription factors were highly represented. Previous data has linked the activity of some ETS family members to HIV expression, and an ETS binding site in the HIV 5’LTR enhancer has been demonstrated (Bassuk et al., 1997; Gegonne et al., 1993; Holzmeister et al., 1993; Leiden et al., 1992; Sieweke et al., 1998). Amongst the negatively associated factors the SNAIL family of transcriptional repressors were identified. No previous reports have connected SNAIL members to HIV. This family of Zn finger TFs has been characterized for their role in regulating epithelial to mesenchymal fate decisions that bind to E-box CANNTG sequences (Nieto, 2002). Notably, the HIV LTR (NL4-3) contains four potential E-box sequences. Additional functional experiments will be needed to confirm whether these novel factors contribute directly or indirectly to HIV gene expression. Through cross-correlation analysis, we have also identified groups of TFs within the HIV-correlated set that are connected to each other, indicating that sets of functionally related TFs can act as part of a coordinated group. As such carefully defining the roles of individual members of these groups will be important.

Notably, the overall correlations of individual TFs and transcripts, although statistically significant, were weak (**Table 1**). We speculate that these low correlations result from the additional influence of background noise or stochastic processes that also contribute to reactivation. Alternatively, this observation could result from latency reversal being determined by the combined activity of several different host cell factors. To address this possibility, we used a machine learning approach to develop a multivariate model to predict whether individual cells expressed viral RNA across the dataset. We successfully developed a model with ~68% accuracy at classifying cells as vRNA+ or vRNA-, confirming that a machine learning approach improves the overall understanding of viral reactivation. It is noteworthy that the most successful model incorporated parameters from all three correlation datasets (correlated transcripts, correlated TF activities, enriched motifs in correlate peaks), although we found that transcript information contributed the most to model performance, followed by TF activity.

To validate this overall approach, we chose to focus on the functional role of one cellular factor, GATA3. Although a previous report has shown that GATA3 can bind an activate an LTR-driven reporter plasmid, the role of GATA3 in HIV transcription and latency is unknown (Yang and Engel, 1993). GATA3 was identified by both correlation of the GATA3 transcript and GATA3 binding motif accessibility with HIV vRNA levels. Across the experimental conditions, GATA3 expression and activity were most clearly induced by prostratin, suggesting a possible role for GATA3 in regulating reactivation of HIV by prostratin. Selective knockdown of GATA3 in 2D10 cells diminished the ability of prostratin to reactivate HIV. Although, clearly, prostratin is not strictly dependent on GATA3 for its latency reversing activity, we propose that GATA3 induction represents one of several pathways activated by prostratin in CD4 T cells that ultimately combine to promote HIV reactivation. As such, a GATA3-targeting drug could be potentially be combined with prostratin or other LRAs to enhance reservoir reactivation during latency reversal therapy. To date, toxicity concerns have limited enthusiasm for PKC agonists such as prostratin in therapy (Jiang and Dandekar, 2015), but a combined stimulation of a PKC agonist with a GATA3 agonist could potentially permit a lower and safer dose of prostratin to be used. Conversely, a GATA3 inhibitor such as pyrrothiogatain could be used to block viral reactivation and promote deeper latency (Nomura et al., 2019). An issue of concern with this approach is that GATA3 plays important physiological roles in promoting the development of important T cell subsets (Zaidan and Ottersbach, 2018). It also remains to be determined whether the role of GATA3 in this response is direct or indirect.

This study should also be considered in light of several inherent limitations and caveats. Firstly, 2D10 cells are an immortalized cell line which have several differences with primary resting CD4 T cells, the natural host cell for the HIV reservoir. Furthermore, the proviral clone contained within these cells is not a full-length HIV strain and has deletions in several viral proteins that could conceivably also contribute to the behavior of the virus in the context of latency reversal (Kim et al., 2011). It is also important to note that our approach rested on examining correlations between cellular factors and viral transcription, with the assumption that some of these correlates represent functional contributors to viral expression. However, it is also possible that while correlated, some factors may be induced by the LRAs tested, but not be causally linked to viral gene reactivation. Other correlated factors may also represent non-contributing “bystanders.” Nevertheless, the confirmation of the functional role of GATA3 in this system demonstrates the power of this approach to identify novel host-viral connections that could be exploited to enhance latency reversal approaches.

## Acknowledgements

This work was supported by the following grants from the National Institutes of Health: NIAID R01 AI143381 (EPB), NIAID UM1 AI164567 (DMM), NIDA R61 DA047023 (EPB), NIAID T32 AI007419 (UNC-Chapel Hill Molecular Biology of Viral Diseases T32, JJP), NIGMS R35 GM138342 (YJ), and NIDA R01 DA054994 (CDR).

## Author contributions

YJ, DM Murdoch and EPB conceived the study. Wet lab experiments were carried out by AM. Computational analysis carried out by JJP, YH and YJ. Machine learning analysis carried out by AO, ZG and CDR. All authors contributed to writing the manuscript.

## Code Availability

Scripts used for analyses carried out in this paper are deposited in the GitHub repository https://github.com/yuchaojiang/HIV-latency.

---

**Supplementary Figure 1.**
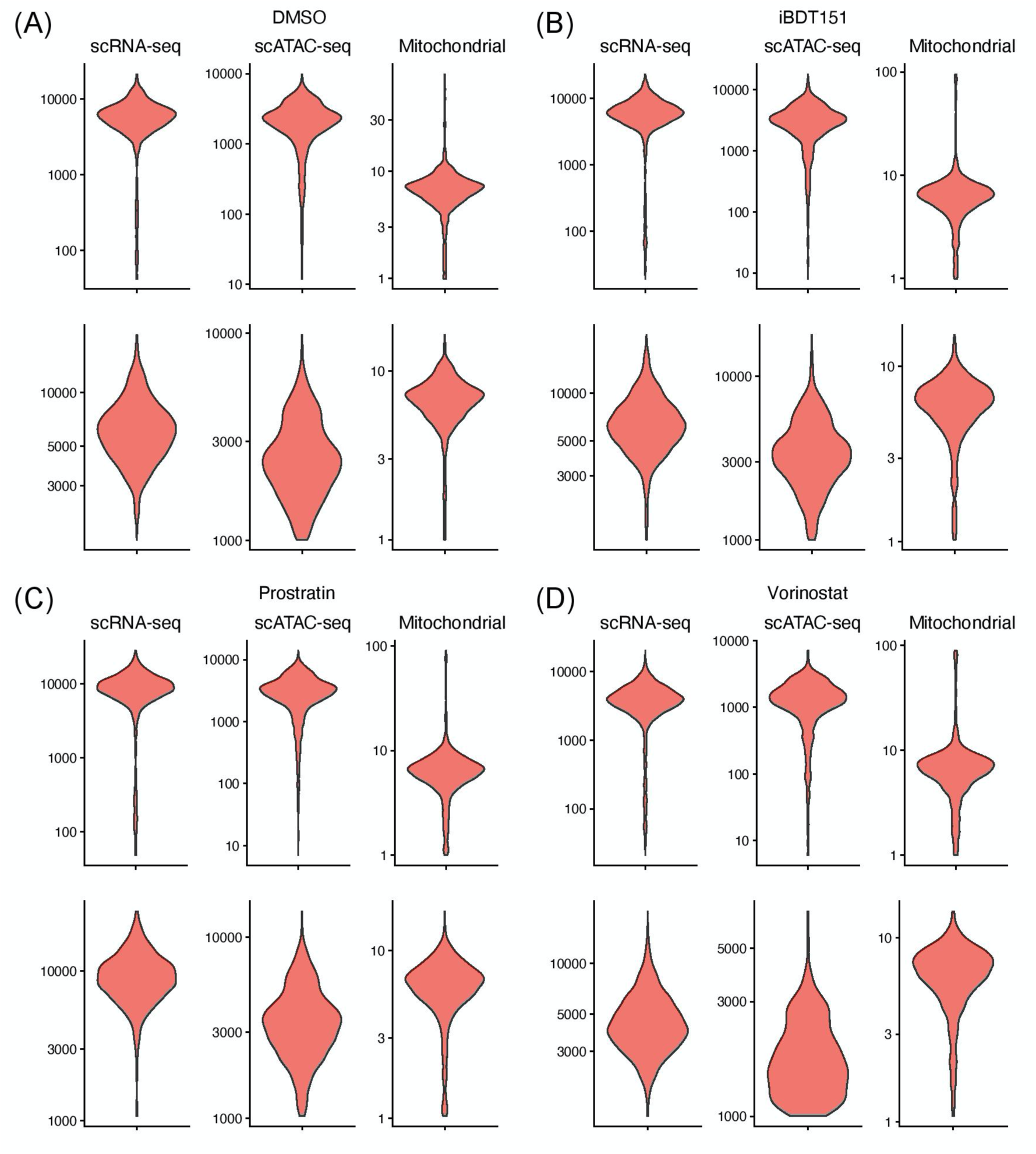
Quality control of the scRNA-seq and scATAC-seq data. **(A-D)** Violin plots showing distribution of scRNA-seq read counts per cell (left), scATAC-seq read counts per cell (middle), and scRNA-seq read counts mapping to the mitochondrial genome (right) for the control sample (DMSO) (A) as well as samples treated with iBET151 (B), prostratin (C), and vorinostat (D). The top and bottom panel correspond to the data before and after filtering based on the quantities.

**Supplementary Figure 2.**
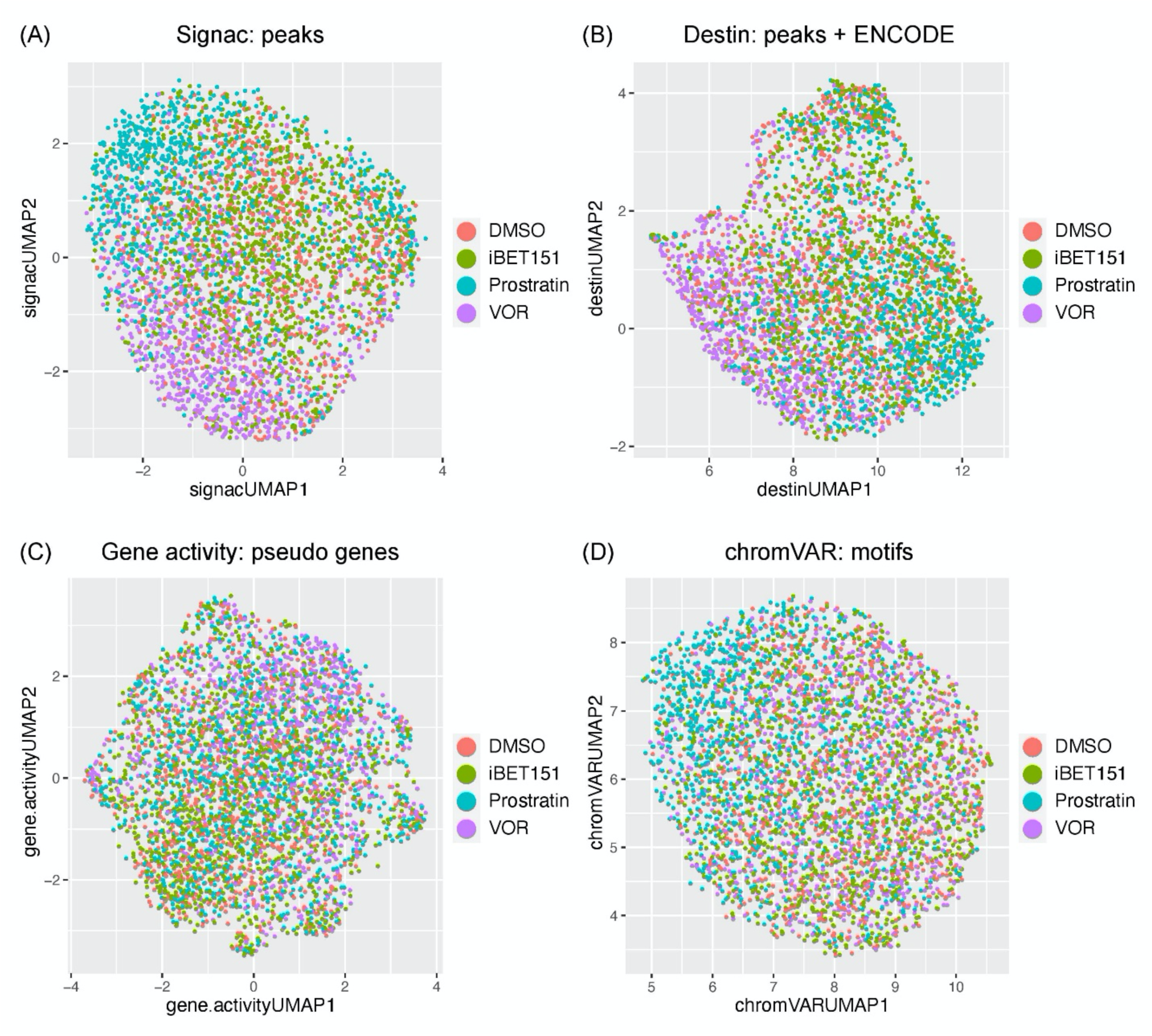
UMAP plots of scATAC-seq data by various methods. **(A-D)** UMAP embeddings by (A) Signac^1^ using the peak count matrix, (B) Destin^2^ using the peak count matrix paired with existing genomic annotations and regulomic data, (C) gene activity^3^ using gene-level measurements by summing up the peak accessibilities in gene bodies and promoter regions, and (D) chromVAR^4^ using motif-level deviation scores.

**Supplementary Figure 3.**
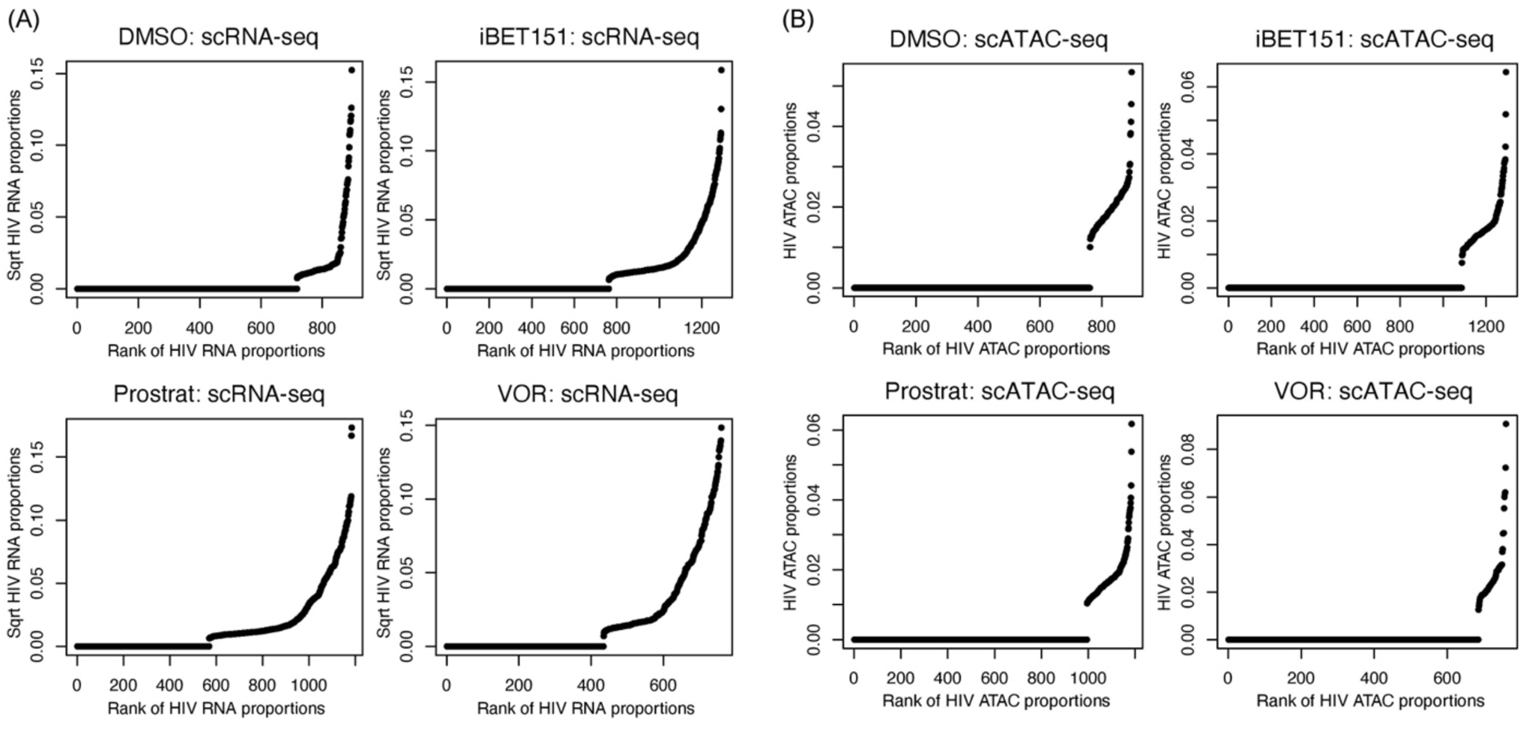
Ranked proportions of the HIV RNA expression and chromatin accessibility across single cells. **(A)** Shown are plots of the square root of the proportions of the scRNA-seq reads mapping to the HIV genes against their ranking across single cells. Each panel is for a different treatment condition. **(B)** Same as (A) but for scATAC-seq reads.

**Supplementary Figure 4.**
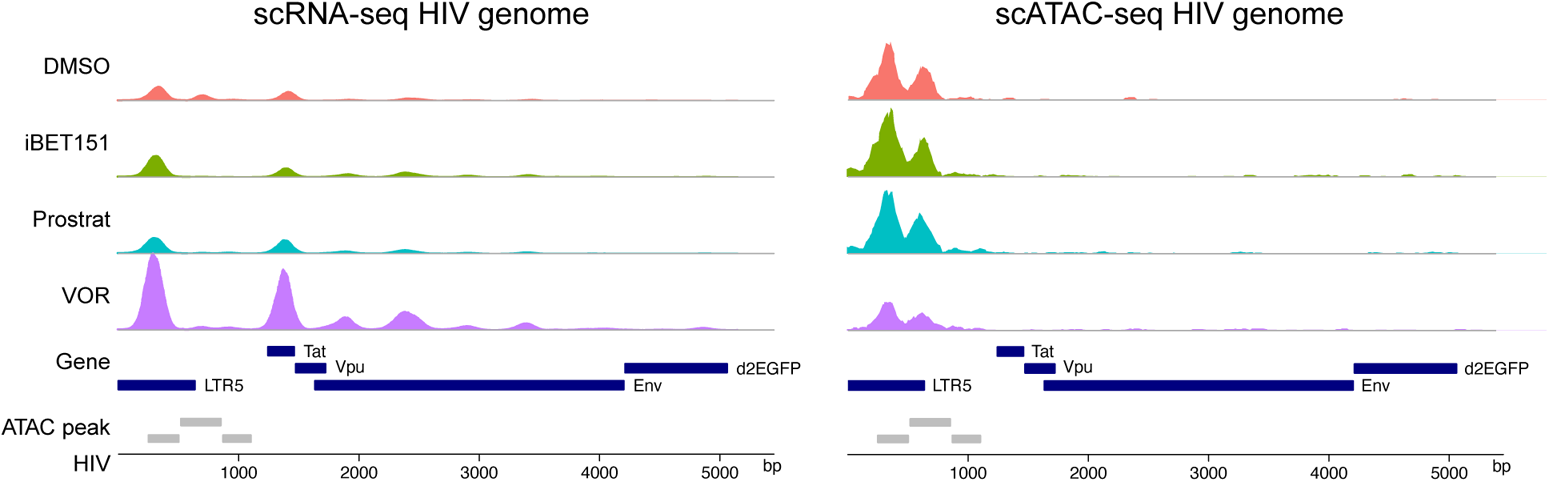
Genome tracks of scRNA-seq and scATAC-seq data per treatment conditions. Coverage plots showing read counts from scRNA-seq (left) and scATAC-seq (right) along the HIV genome for different treatment conditions. The counts were normalized against the cell number within each condition. The dark blue and gray boxes represent gene annotations and accessible chromatin regions based on ATAC-seq signals (ATAC peak).

**Supplementary Figure 5.**
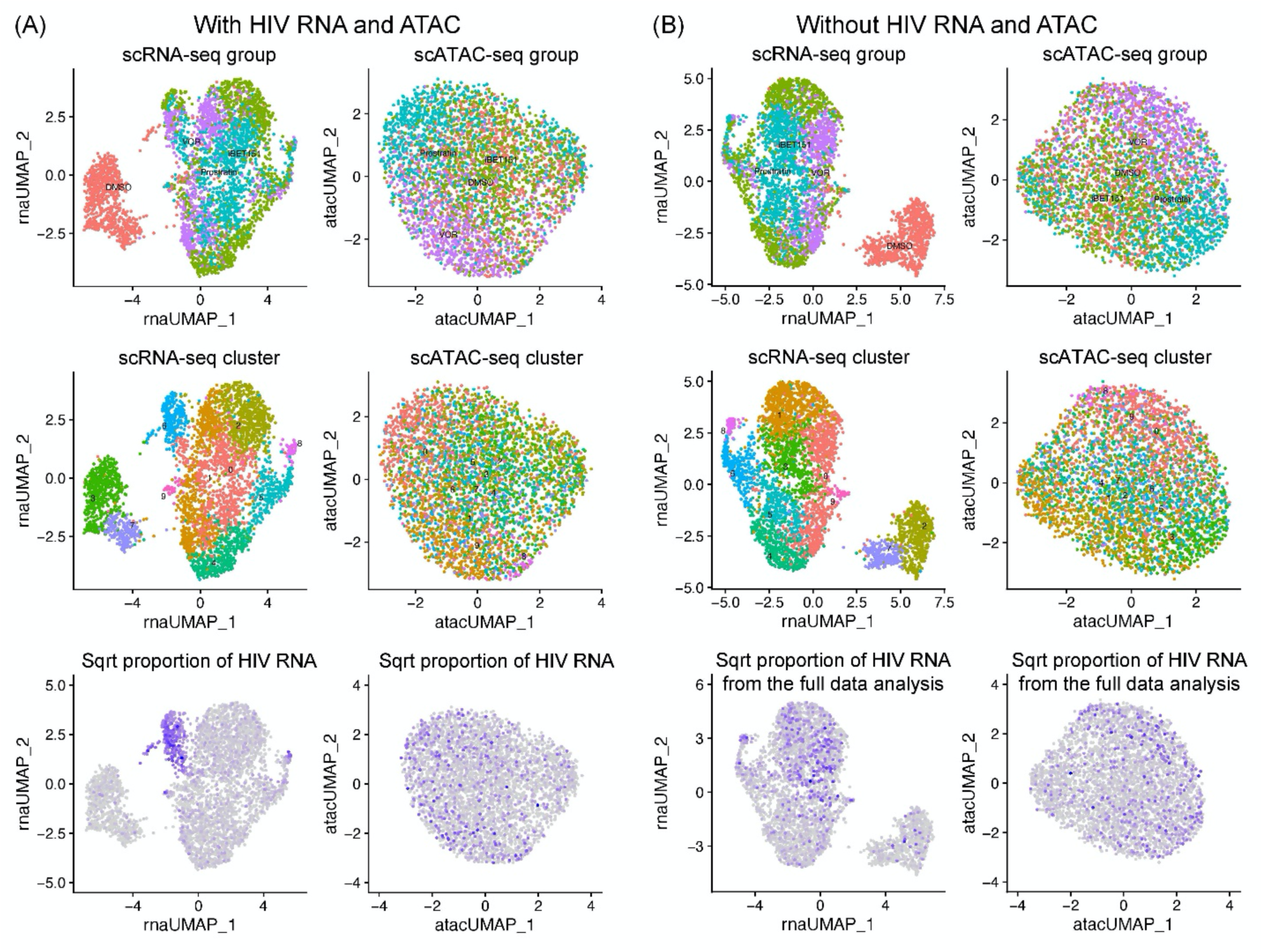
Cluster 6 is driven by HIV RNA expression. **(A)** UMAP plots of scRNA-seq (left) and scATAC-seq (right) data. The colors represent treatment conditions (top), clusters (middle), and proportions of the HIV RNA expression from the full data analysis (bottom). **(B)** Same as (A) except that the HIV genes were excluded for the clustering and UMAP construction.

**Supplementary Figure 6.**
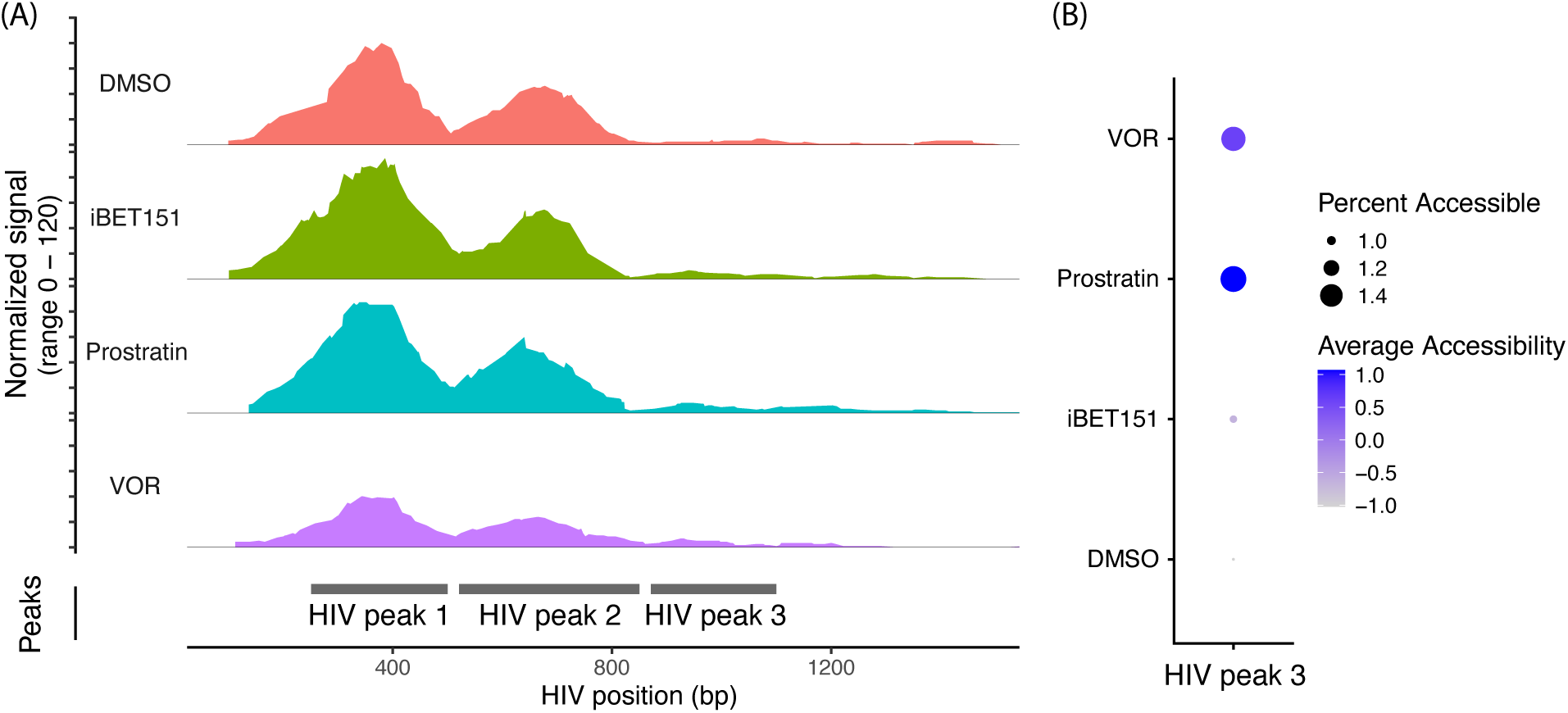
ATAC-seq signals within three chromosomal regions in the HIV genome. **(A)** Coverage plots showing read counts from scRNA-seq in the 5’ region of the HIV genome per treatment conditions. The gray boxes represent ATAC peak regions. **(B)** Dot plots showing chromatin accessibility in the third peak (HIV peak 3) for different treatment conditions.

**Supplementary Figure 7.**
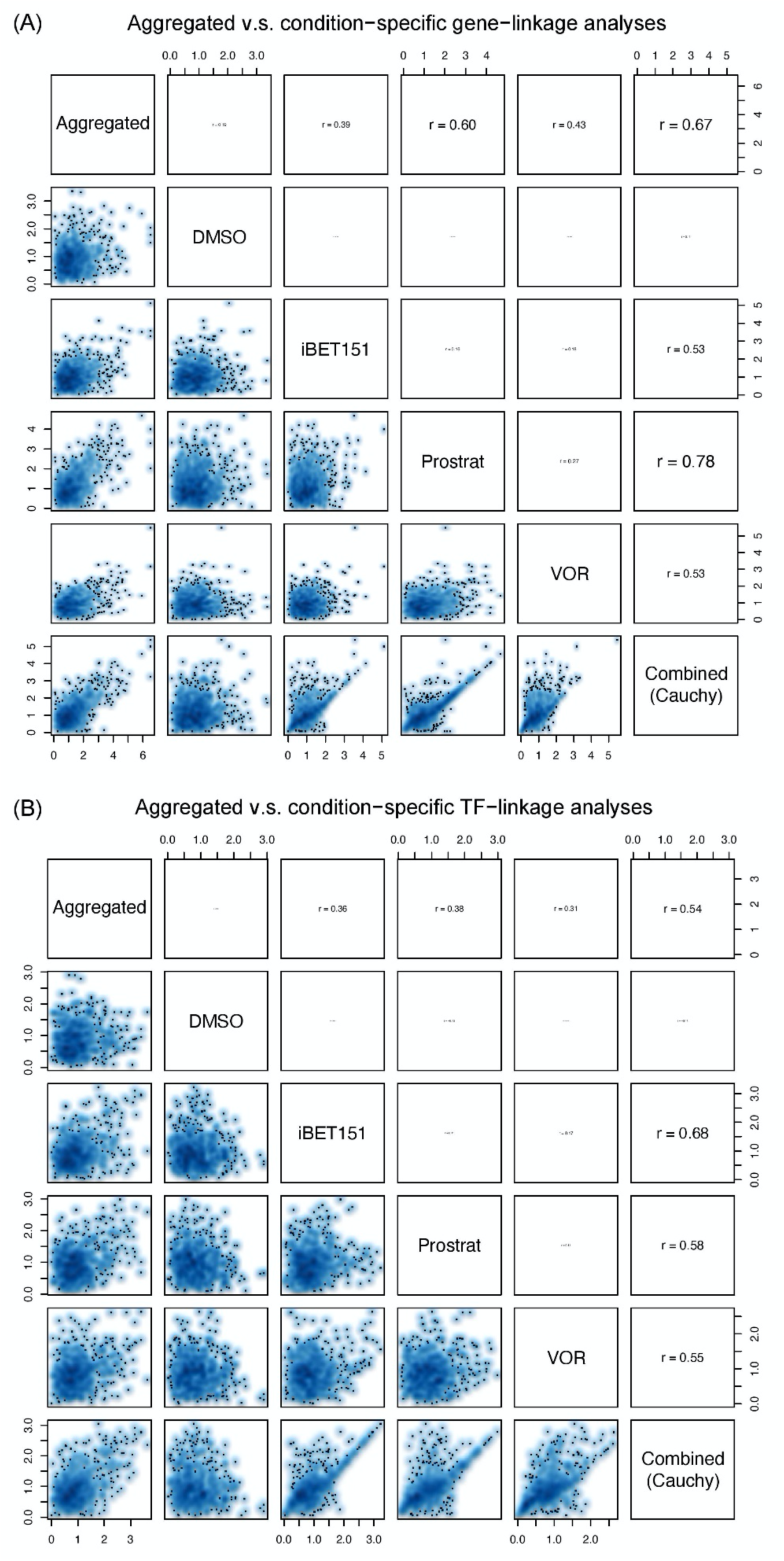
Comparison of gene/TF-linkage analysis between the aggregated and condition-specific samples. **(A)** Pairwise scatterplots comparing the square root of negative log-transformed nominal *p*-values from tests for a non-zero slope in simple linear regression of the HIV expression on cellular gene expression between different treatment conditions. The combined *p*-values were obtained from the samples treated with iBET151, prostratin (Prostrat), and VOR. **(B)** Same as (A) but for TF motif scores.

**Supplementary Figure 8.**
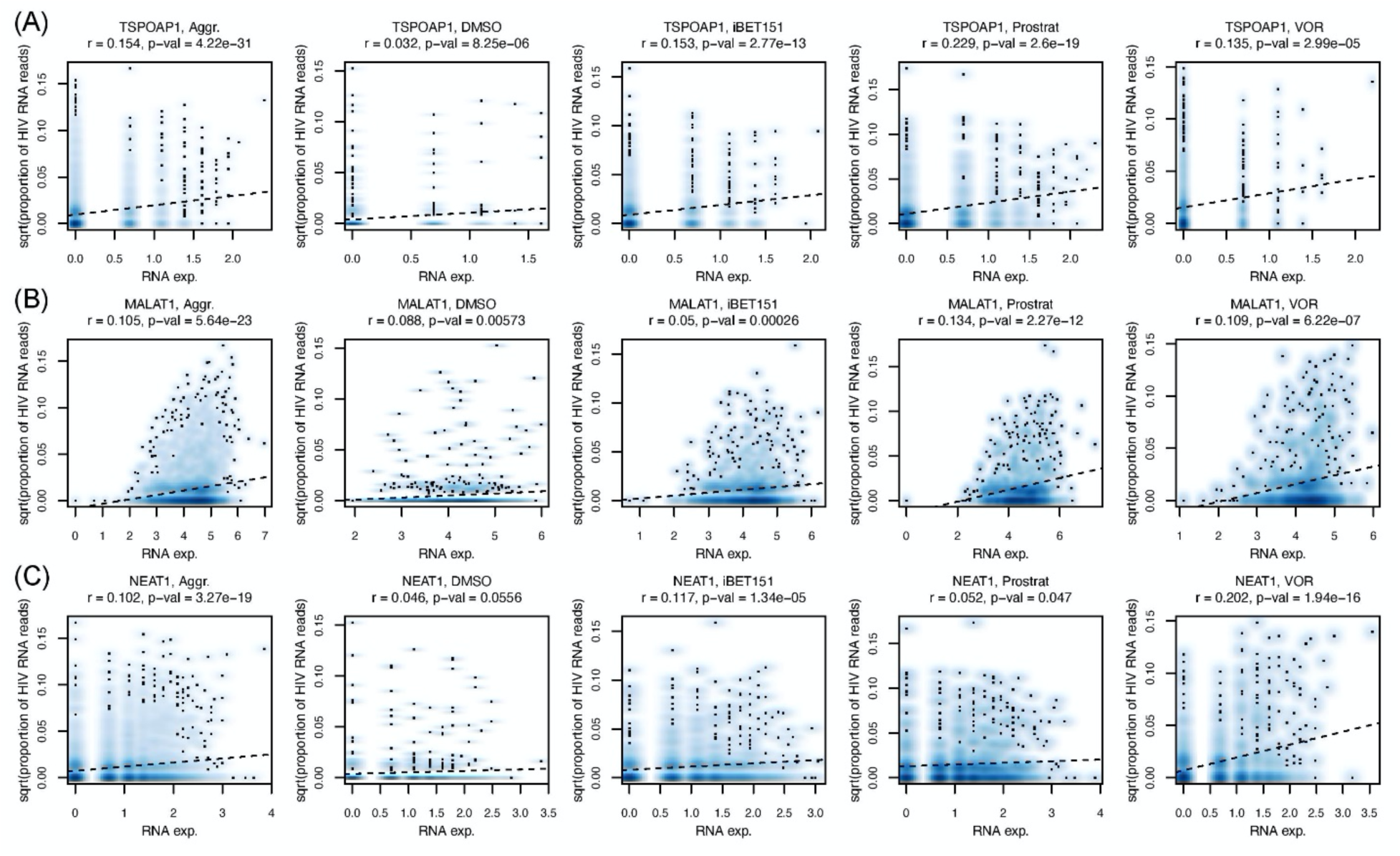
Representative association patterns between HIV RNA expression and cellular gene expression from aggregated and condition-specific analyses. **(A)** Scatter plots comparing the square root of the proportion of the HIV RNA expression and the expression level of the *TSPOAP1* gene. Each point represents a single cell. Spearman correlation coefficients and nominal *p*-values from tests for non-zero slope in simple linear regression are also shown. The dotted line represents the fitted line from regression. **(B)** Same as (A) but for *MALAT1*. **(C)** Same as (A) but for *NEAT1*.

**Supplementary Figure 9.**
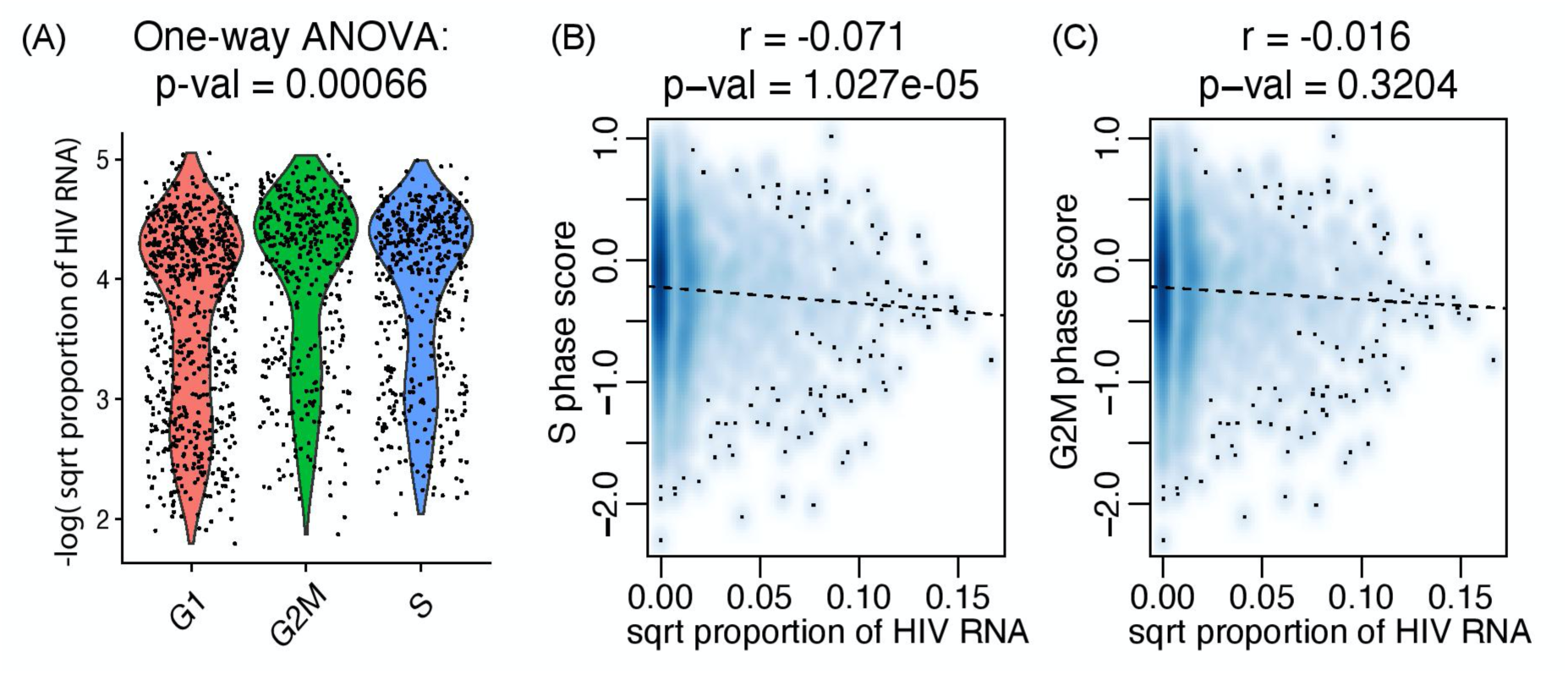
HIV RNA expression is negatively associated with S phase. **(A)** Violin plots showing distribution of G1-, G2M-, and S-phase scores across single cells. Shown on the top is a *p*-value from a one-way ANOVA test comparing the means. **(B)** A scatter plot comparing the square root of proportion of HIV RNA reads against the S phase scores. Each point represents a single cell. Spearman correlation (r) and associated *p*-value (p) are also shown. **(C)** The same as in (B) but for the G2M scores.

**Supplementary Figure 10.**
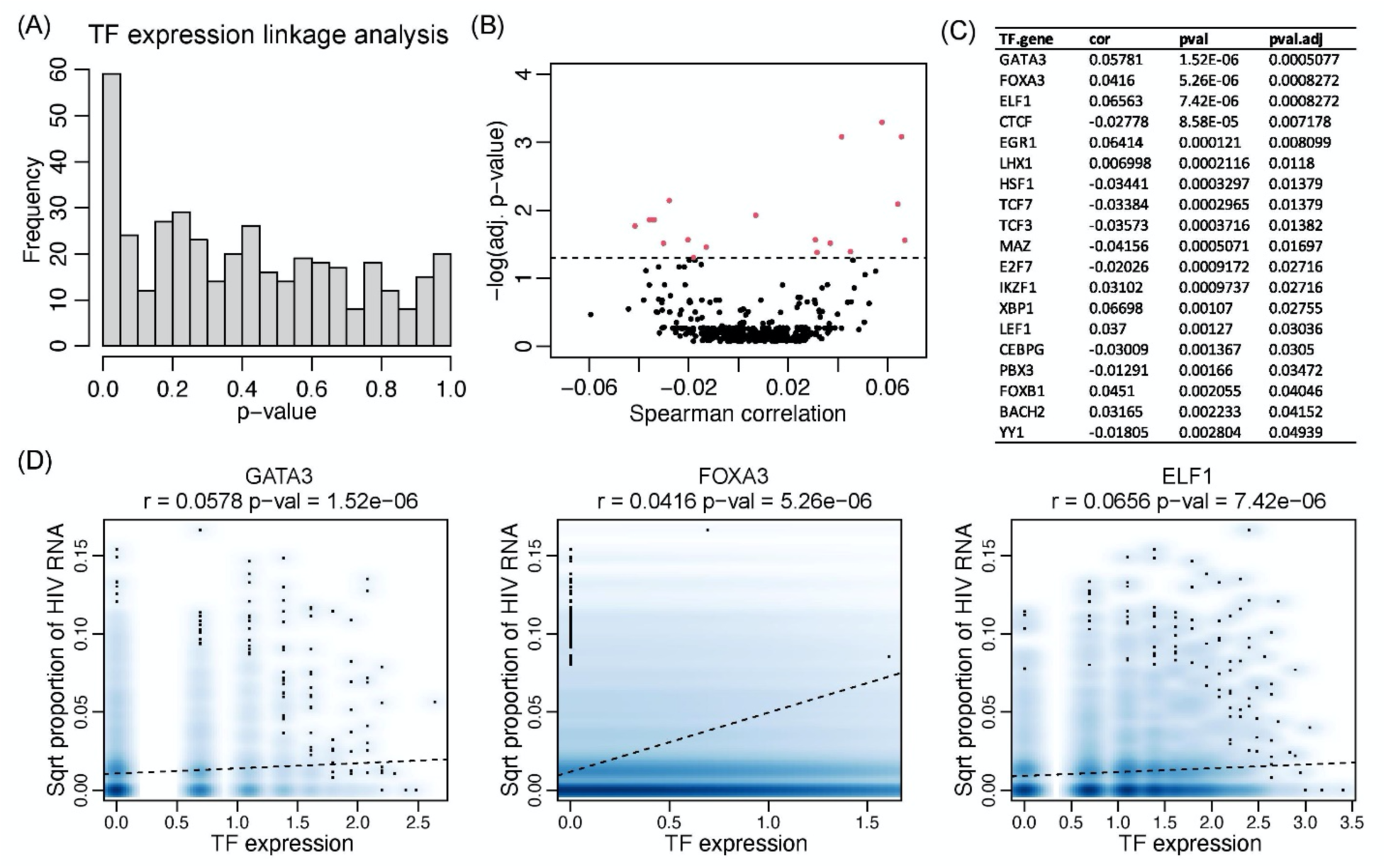
Associations between HIV RNA expression and genes encoding TFs. **(A)** A histogram showing distribution of nominal *p*-values from tests for a non-zero slope in simple linear regression of the HIV RNA expression on TF gene expression. **(B)** Shown is a plot of the transformed adjusted *p*-values from simple linear regression as in (A) against Spearman correlation coefficients between the HIV RNA expression and TF gene expression. Each point represents a gene encoding a TF. The dotted line represents the adjusted *p*-value of 0.05. Points with *p*-values below this threshold are colored in red. **(C)** Shown are Spearman correlation coefficients (cor), nominal *p*-values from linear regression, and adjusted *p*-values (pval.adj) for top-ranked genes encoding TFs. **(D)** Scatter plots comparing the square root of the proportion of the HIV RNA expression and the expression levels of genes encoding the GATA3 (left), FOXA3 (middle), and ELF1 (right) TFs. Spearman correlation coefficients and nominal *p*-values from tests for non-zero slope in simple linear regression are also shown. The dotted line represents the fitted line from regression.

**Supplementary Figure 11.**
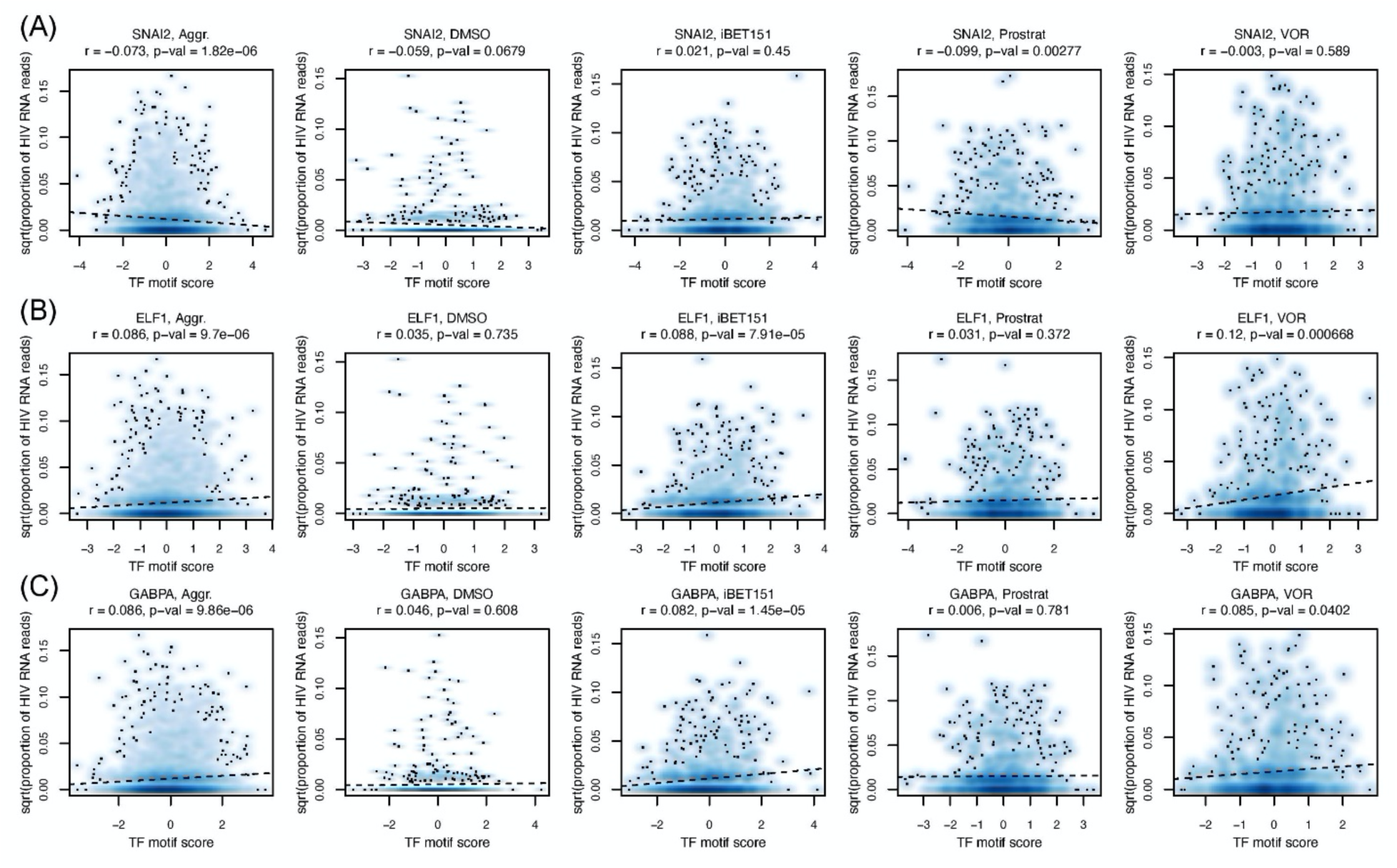
Representative association patterns between HIV RNA expression and TF motif scores from aggregated and condition-specific analyses. **(A)** Scatter plots comparing the square root of the proportion of the HIV RNA expression and the TF motif scores for the SNAI2 TF. Each point represents a single cell. Spearman correlation coefficient and nominal *p*-value from a test for non-zero slope in simple linear regression are also shown. The dotted line represents the fitted line from regression. **(B)** Same as (A) but for the ELF1 TF. **(C)** Same as (A) but for the GABPA TF.

**Supplementary Table 1.** Genes that were significantly up-regulated and down-regulated upon treatment with LRAs. The columns contain gene names (gene), nominal *p*-values from Wilcoxon rank sum tests comparing expression levels between the control (DMSO) and one of the LRA-treated samples (p_val), log fold changes of the average expression between the control (DMSO) and LRA-treated samples (avg_log2FC), the percentage of cells where the gene is detected in the LRA-treated sample (pct.1), the percentage of cells where the gene is detected in the control sample (pct.2), and adjusted *p*-values (p_val_adj), as implemented in Seurat (ref). Results are separately attached as an Excel spreadsheet.

**Supplementary Table 2.** Gene ontology (GO) term enrichment analyses for gene that were significantly up-regulated and down-regulated upon treatment with LRAs. Analyses of GO enrichment (biological process) were carried out for each condition using DAVID^5^. Results are separately attached as an Excel spreadsheet.

**Supplementary Table 3.** Identification of cellular transcripts that correlate with HIV RNA levels. FDR adjusted *p*-values are included. Results are separately attached as an Excel spreadsheet.

**Supplementary Table 4.**
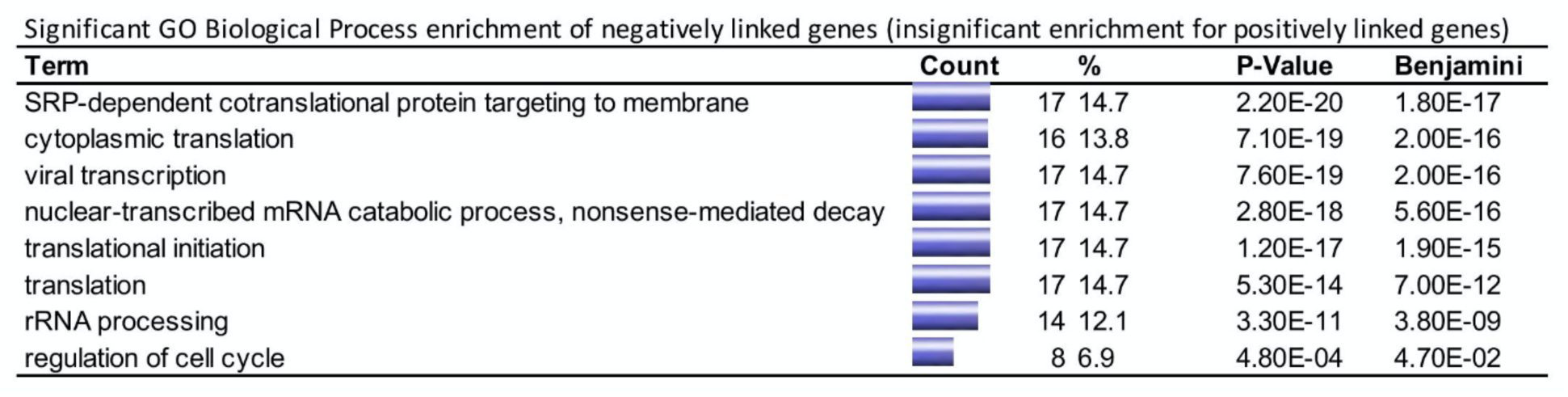
GO “biological process” enrichment analysis using genes that were significantly linked with HIV RNA expression. Only negatively linked genes led to significant enrichments (insignificant enrichment for positively linked genes). The columns contain counts of the associated genes in the GO categories, percentages of the associated genes in the GO categories, nominal *p*-values from hypergeometric tests, and Benjamini-Hochberg-corrected *p*-values.

**Supplementary Table 5.** Identification of ATAC peak regions that correlate with HIV RNA levels. The column contains ATAC peak regions (peak), Spearman correlation between chromatin accessibility in the peak regions and HIV RNA levels (cor), and nominal *p*-values from tests for non-zero slope in simple linear regression (pval). Results are separately attached as an Excel spreadsheet.

**Supplementary Table 6.**
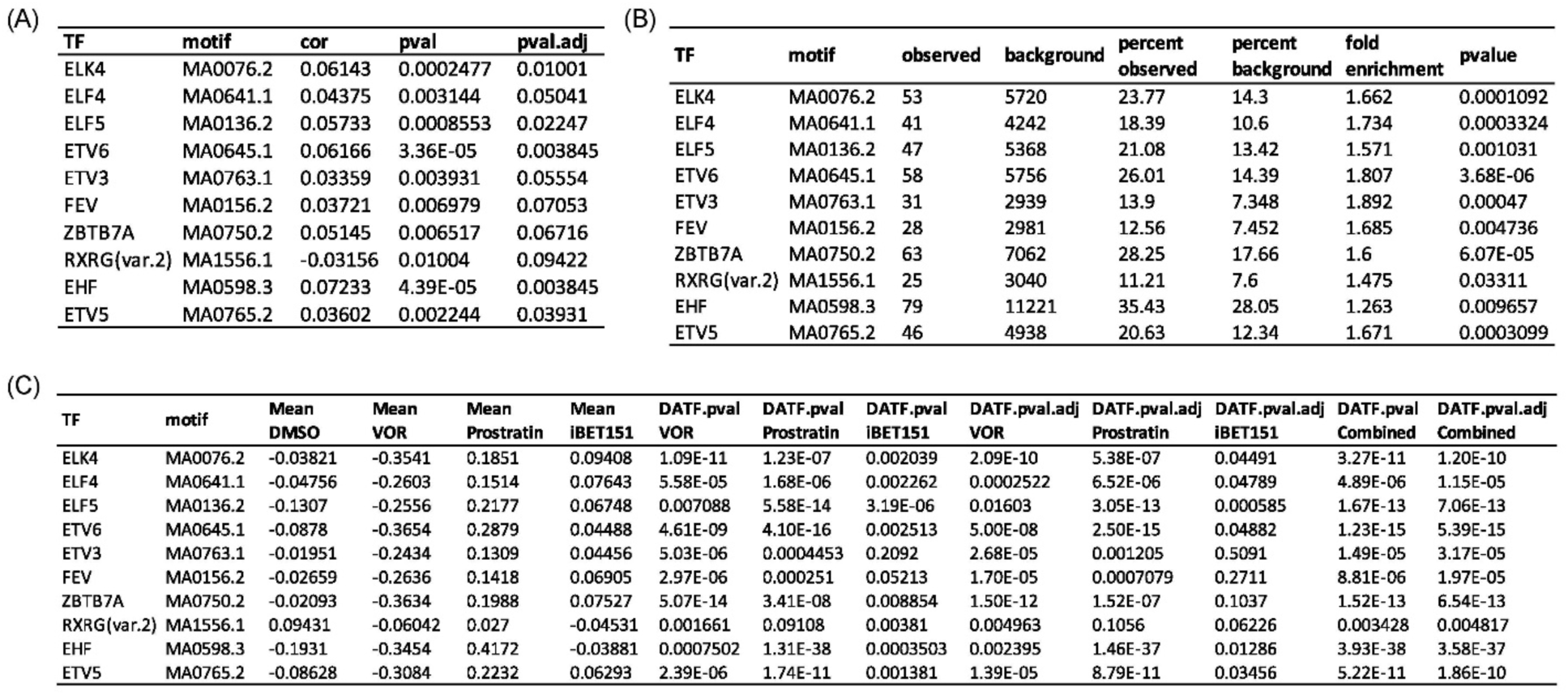
Testing statistics for ten TFs that are significant from three testing schemes. Ten TFs (Figure 6A) remain significant from the TF linkage analysis, enrichment analysis of TF-binding motifs using significantly linked peaks, and analysis of differential motif accessibility. **(A)** Associations between HIV RNA expression and TF motif scores. See the legend to Table 1B. **(B)** Motifs enrichment analysis. See the legend to Table 1D. **(C)** Differential accessibility analysis upon LRA treatment. The table contains TF names, motif names, the mean of chromatin accessibility for different treatment conditions, nominal *p*-values from Wilcoxon rank sum tests comparing chromatin accessibility in the motifs between the control (DMSO) and one of the LRA-treated samples (p_val), adjusted *p*-values (p_val_adj), and combined *p*-values.

**Supplementary Table 7.** Master output of TF-specific testing. Testing results from the TF linkage analysis (linkedTF), enrichment analysis of TF-binding motifs using significantly linked peaks (enrichedTF), and analysis of differential motif accessibility (DATF) across all TFs. *P*-values are integrated using the Cauchy combination test.

